# A global resource for genomic predictions of antimicrobial resistance and surveillance of *Salmonella* Typhi at Pathogenwatch

**DOI:** 10.1101/2020.07.03.186692

**Authors:** Silvia Argimón, Corin A. Yeats, Richard J. Goater, Khalil Abudahab, Benjamin Taylor, Anthony Underwood, Leonor Sánchez-Busó, Vanessa K. Wong, Zoe A. Dyson, Satheesh Nair, Se Eun Park, Florian Marks, Andrew J. Page, Jacqueline A. Keane, Stephen Baker, Kathryn E. Holt, Gordon Dougan, David M. Aanensen

## Abstract

**Background:** Microbial whole-genome sequencing (WGS) is now increasingly used to inform public health investigations of infectious disease. This approach has transformed our understanding of the global population structure of *Salmonella enterica* serovar Typhi (*S.* Typhi), the causative agent of typhoid fever. WGS has been particularly informative for understanding the global spread of multi-drug resistant (MDR) typhoid. As WGS capacity becomes more decentralised, there is a growing opportunity for collaboration and sharing of surveillance data within and between countries to inform disease control policies. This requires freely available, community driven tools that reduce the barriers to access genomic data for public health surveillance and that deliver genomic data on a global scale.

**Methods:** Here we present the Pathogenwatch (https://pathogen.watch/styphi) scheme for *S.* Typhi, a web application enabling the rapid identification of genomic markers of antimicrobial resistance (AMR) and contextualization with public genomic data to identify high-risk clones at a population level. Data are delivered in single genome reports or in collections of genomes combined with geographic and other data using trees, maps and tables.

**Results:** We show that the clustering of *S.* Typhi genomes in Pathogenwatch is comparable to established bioinformatics methods, and that genomic predictions of AMR are largely concordant with phenotypic drug susceptibility data. We demonstrate the public health utility of Pathogenwatch with examples selected from over 4,300 public genomes available in the application.

**Conclusions:** Pathogenwatch democratises genomic epidemiology of *S.* Typhi by providing an intuitive entry point for the analysis of WGS and linked epidemiological data, enabling international public health monitoring of the emergence and spread of high risk clones.

## Introduction

Bacterial pathogens have the potential for rapid evolution and adaptation (1). The ability to rapidly sequence microbial genomes directly from the field allows tracking of pathogen evolution in real-time and in a geographical context. Genomic surveillance provides the opportunity to identify the emergence of genetic signatures indicating antimicrobial resistance (AMR), or adaptation to host, facilitating early intervention and minimising wider dissemination. Consequently, genomic data has the ability to transform the way in which we manage the emergence of microbes that pose a direct threat to human health in real time.

Although pathogen genomic data is being generated at a remarkable rate, we need to bridge the gap between genome sciences and public health with tools that make these data broadly and rapidly accessible to those who are not expert in genomics. To maximise the impact of ongoing surveillance programs, these tools need to quickly highlight high-risk clones by assigning isolates to distinct lineages and identifying genetic elements associated with clinically relevant features such as AMR or virulence. In this way, new isolates can be examined against the backdrop of a population framework that is continuously updated and that enables both the contextualisation of local outbreaks and the interpretation of global patterns.

*Salmonella enterica* subsp. *enterica* serovar Typhi (*S.* Typhi) causes typhoid (enteric) fever, a disease that affects approximately 20-30 million people every year (2, 3). The disease is predominant in low-income communities where public health infrastructure is poorly resourced. Similar to other infections, typhoid treatment is compromised by the emergence of *S.* Typhi with resistance to multiple antimicrobials, including those currently used for treatment (3). Until recently, epidemiological investigations and surveillance of typhoid fever have employed alternative molecular techniques such as pulse-field gel electrophoresis (PFGE (4)), multi-locus sequence typing (MLST (5)), multiple-locus variable-number tandem-repeat (VNTR) analysis (MLVA (6)), and phage-typing (4), which offer insufficient resolution for a bacterium that exhibits very limited genetic variability. Whole genome sequencing (WGS) has proven key to identify *S.* Typhi high-risk clones by linking the population structure to the presence of AMR elements. For example, the recent resurgence of multi-drug resistant (MDR) typhoid (defined as resistance to all the historical first-line agents chloramphenicol, ampicillin and co-trimoxazole) has been explained in part by the global spread of an MDR *S*. Typhi lineage known as haplotype H58 or subclade 4.3.1 (7, 8), which is associated with both acquired AMR genes (conferring MDR) and fluoroquinolone resistance mutations (7, 9).

WGS is increasingly being implemented in local and national public health laboratories, and web applications can provide rapid analysis and access to actionable information for infection control in the context of a global population framework. Online resources are available for the identification of acquired AMR mechanisms in bacterial pathogens, including *Salmonella spp.* (10, 11), and for *in silico* typing and visualisation of genome variation and relatedness based on WGS data (12–16), albeit without an emphasis on typhoidal *Salmonella*. Here, we describe Typhi Pathogenwatch, a web application to support genomic epidemiology and public health surveillance of *S.* Typhi. Typhi Pathogenwatch rapidly places new genomes within the broader geographic and population context, predicts their genotype according to established nomenclatures (5, 8, 12), and detects the presence of AMR determinants and plasmid replicon genes to assess public health risk. Typhi Pathogenwatch displays this information interactively, allowing users to link lineages, AMR profiles, geographical data and other metadata to quickly determine if similar strains have been previously identified, where and when. Furthermore, results can be downloaded or shared via a web address containing a unique collection identifier. This approach allows the rapid incremental addition of new data and can be used to underpin the international surveillance of typhoid, MDR and other public health threats.

## Methods

### The Pathogenwatch application

The Pathogenwatch user interface is a React (17) single-page application with styling based on Material Design Lite (18). Phylocanvas (19) is used for phylogenetic trees, Leaflet (20) is used for maps, and Sigma (21) is used for networks. The Pathogenwatch back-end, written in Node.js, consists of an API service for the user interface and four “Runner” services to perform analysis: species prediction, single-genome analyses, tree-building, and core genome multi-locus sequence typing (cgMLST) clustering. Runner services spawn Docker containers for queued tasks, streaming a FASTA file or prior analysis through standard input and storing JSON data from standard output. Data storage and task queuing/synchronisation are performed by a MongoDB cluster.

### *S.* Typhi genome assemblies and data privacy

Genome assemblies can be uploaded by the user in FASTA format or assembled *de novo* from high-throughput short read data with the Pathogenwatch pipeline using SPAdes (22), as described in the Pathogenwatch documentation (23). Sequence data and metadata files uploaded by the user are kept private to the user account unless explicitly requested to be publicly shared. Genomes can be grouped into collections and kept private or set to be made available to collaborators through a web link. Users can also integrate private and potentially confidential metadata into the display without uploading it to the Pathogenwatch servers. This private metadata will not be shared even if the collection is set to be shared via web link (24).

Genomes from published studies with geographical localisation metadata and short read data on the European Nucleotide Archive (ENA) are available as public data and accessible to all users for browsing and for contextualisation of their own datasets. As of November 2020, 4389 public *S.* Typhi genomes from 26 studies were available (Additional File 1: Supplementary Table S1), either sequenced and available from the Wellcome Sanger Institute (WSI) directly, or obtained from the ENA. Genomes sequenced at WSI were assembled *de novo* with a previously described assembly pipeline (25). Briefly, FASTQ files were used to create multiple assemblies using VelvetOptimiser v2.2.5 and Velvet v1.2 (26). An assembly improvement step was applied to the assembly with the best N50, and contigs were scaffolded using SSPACE (27) and sequence gaps filled using GapFiller (28). Genomes downloaded from the ENA were assembled with Velvet as above, as well as with SPAdes v3.9.0 (22) and a range of *k*-mer sizes of 66-90% of the read length (in increments of 4). Assemblies were evaluated based on their metrics and the Pathogenwatch core genome stats (number of contigs, assembly length, N50, non-ATCG characters, GC content, number of core matches,). Seventeen public and published genomes were excluded as the assemblies either contained more than 700 contigs, more than 50,000 non-ATCG characters, a GC content below the smallest GC content or above than the largest GC content of the *S. enterica* subsp *enterica* genomes in RefSeq. or a total length that is <10% smaller than the smallest genome or >10% larger than the largest *S. enterica* subsp *enterica* genome in RefSeq, For five isolates, we used genome assemblies deposited in GenBank that met the same quality criteria. The metadata and assembly stats and method of the public genomes is available on (Additional File 2: Supplementary Table S2).

### Characterisation and genotyping of *S.* Typhi genomes with Pathogenwatch

For both user-uploaded and public genomes, Pathogenwatch outputs a taxonomy assignment, a map of their locations, and assembly quality metrics. The taxonomy assignment is the best match to a microbial version of the RefSeq genome database release 78, as computed with Mash (29) (k=21, s=400). Details of the *speciator* tool can be found in the documentation (30).

Pathogenwatch also provides compatibility with *Salmonella* serotyping (SISTR (15)), multi-locus sequence typing (MLST (5)), core-genome MLST (cgMLST (12)) and *S.* Typhi single-nucleotide polymorphism (SNP)-based genotyping (GenoTyphi (8)). Detailed descriptions of the implementation of the typing tools can be found in the documentation (31).

The MLST and cgMLST schemes are periodically downloaded from Enterobase (32) and (33), respectively. Samples are typed as described in the documentation (https://cgps.gitbook.io/pathogenwatch/technical-descriptions/typing-methods/mlst and https://cgps.gitbook.io/pathogenwatch/technical-descriptions/typing-methods/cgmlst). Exact allele matches are reported using their allele ID. Multiple allele hits for a gene are reported if present. Inexact allele matches and novel STs are reported by hashing the matching allele sequence and the gene IDs, respectively.

Pathogenwatch implements SISTR (Salmonella In Silico Typing Resource (15)), which produces serovar predictions from WGS assemblies by determination of antigen gene and cgMLST gene alleles using *blastn* v2.2.31+. Pathogenwatch uses the cgmlst_subspecies and serovar fields from the SISTR JSON output to specify the serovar.

GenoTyphi assigns *S.* Typhi genomes to a predefined set of clades and subclades based on a curated set of SNPs (8) that is regularly updated as novel lineages of epidemiological interest are identified (34). Pathogenwatch uses an in-house implementation designed to work with assembly output. The *blastn* v2.2.30 program is used to align the query loci and identify positions of diagnostic SNPs, which are then processed according to the rules of the GenoTyphi scheme (35). The genotype assignment and the number of diagnostic SNPs identified on the assemblies are reported.

The plasmid replicon marker sequences are detected in the user and public genome assemblies with *Inctyper*, which uses the PlasmidFinder Enterobacteriaceae database (36). Details of the *Inctyper* tool can be found in the documentation (37).

### The Pathogenwatch *S.* Typhi core genome library

Pathogenwatch supports SNP-based neighbour joining trees of *S.* Typhi both for user genomes (collection trees) and public genomes (population tree and subtrees). The trees are inferred using a curated core gene library of 3284 *S.* Typhi genes (38) generated from a pan-genome analysis of 26 complete or high-quality draft genomes (Additional File 1: Supplementary Table S3) with Roary (39) and identity threshold of 95%. The core gene families were realigned using MAFFT v7.2.2.0 (40), and filtered or trimmed according to the quality of the alignments. The gene with the fewest average pairwise SNP differences to the other family members was selected as the representative for each family. We then selected 19 reference genomes (Additional File 1: Supplementary Table S3) belonging to different genotypes according to the population structure previously described (8). The gene families were then searched against each of the 19 reference genomes and filtered according to the following rules: a) only universal families with complete coverage of the representative were kept; b) all paralogues were removed; c) overlapping gene families were merged into a single, contiguous pseudo-sequence. A BLAST (41) core library was then built with the representative genes, and a profile of variant sites determined for the core genes present in each reference genome. Each of the 4389 public genomes was then clustered with its closest reference genome based on this profile of variant sites, thus constituting each of the 19 population subtrees that Pathogenwatch employs to contextualise user-uploaded genomes.

### Pathogenwatch genome clustering of *S.* Typhi

The relationships between genomes are represented with trees (dendrograms) based on the genetic distance computed from substitution mutations in the core gene library, as described in detail in the documentation (42). User-provided assemblies are queried against the *S.* Typhi core gene library with *blastn* v2.2.30 (41) using an identity threshold of 90%. The core gene set of each query assembly is compared to the reference genome core that has the most variant sites in common. An overall relative substitution rate is determined, and loci that contain more variants than expected assuming a Poisson distribution are filtered out. Pairwise distances between assemblies (including user-provided and reference) are scored via a distance scoring algorithm that compares all variant positions from all pairs of core gene sets, SNPs are counted (generating a downloadable pairwise difference matrix) and normalised by the relative proportion of the core present (generating a downloadable pairwise score matrix). The pairwise score matrix is then used to infer a midpoint-rooted neighbour-joining tree using the Phangorn v2.4.0 (43) and Ape v5.1 (44) R packages. Trees are computed for the user assemblies only (collection tree), and for the user assemblies and public assemblies assigned to the same reference genome (public data subtrees), all of which are downloadable in Newick format.

We benchmarked the Pathogenwatch clustering method against other methods of SNP-based tree inference with three subsets of published genomes: Dataset I) 118 genomes spanning the population diversity of *S.* Typhi defined by GenoTyphi (Additional File 3: Supplementary Table S4); Dataset II) 138 closely related genomes, from a recent clonal expansion of the multidrug-resistant haplotype H58 within Africa (Additional File 2: Supplementary Table S5); and Dataset III) 43 strains from clade 3.2 including CT18, the first completed *S.* Typhi genome, which remains reference of choice for most population genomics studies (Additional File 2: Supplementary Table S6). For each subset a tree was generated with four different methods: 1) Pathogenwatch; 2) maximum likelihood (ML) with RAxML v8.2.8 (45) on SNPs extracted from an alignment of concatenated core genes generated using Roary (39); 3) neighbour joining (NJ) with FastTree (46) using the option –noml on the same alignment as 2); and 4) ML with RAxML v8.2.8 on SNPs extracted from a previously published CT18-guided alignment (7). Five hundred bootstrap replicates were computed for the ML trees (methods 2 and 4). We compared the trees thus generated using the tree comparison software Treescape v1.10.18 (Kendall-Colijn distance, now available as Treespace (47)) and the Tanglegram algorithm of Dendroscope (48). The tree files used in the tree comparisons are provided in (49).

Genomes can also be clustered in Typhi Pathogenwatch based on their cgMLST profile using single linkage clustering. Distance scores are calculated between each pair of samples by identifying the genes which have been found in both samples and by counting the number of differences in the alleles. The SLINK algorithm (50) is used to quickly group genomes into clusters at a given threshold. For a given genome, users are able to see how many other genomes it is clustered with at a range of distance thresholds, view the structure of the cluster as a network graph, and view the metadata and analysis for sequences in that cluster.

### Genomic predictions of antimicrobial resistance

The Pathogenwatch AMR prediction module queries the genome assemblies with *blastn* v2.2.30 (41) for the presence of genes and single point mutations known to confer resistance in *S.* Typhi to ampicillin (AMP), chloramphenicol (CHL), broad-spectrum cephalosporins (CEP), ciprofloxacin (CIP), sulfamethoxazole (SMX), trimethoprim (TMP), the combination antibiotic co-trimoxazole (sulfamethoxazole-trimethoprim, SXT), tetracycline (TCY), azithromycin (AZM), colistin (CST) and meropenem (MEM) (Additional File 1: Supplementary Table S7 (51)). For details of the implementation see the Pathogenwatch documentation (52).

The Pathogenwatch AMR prediction module also provides a prediction of AMR phenotype inferred from the combination of identified mechanisms. To benchmark the genotypic resistance predictions, we used a set of 1316 genomes from 16 published studies (Additional File 1: Supplementary Table S1) with drug susceptibility information available for at least one of the twelve antibiotics reported by Typhi Pathogenwatch. The drug susceptibility data reported was heterogeneous across the studies (minimum inhibitory concentration (MICs), disk diffusion diameters, and/or susceptible/intermediate/resistant (SIR)). We first compared the Typhi Pathogenwatch antibiotic resistance predictions to the drug susceptibility phenotype (SIR interpretation provided by the studies) of 1316 genomes, grouping the Resistant and Intermediate classifications as non-susceptible. For each antibiotic, the sensitivity, specificity, positive predictive value (PPV) and negative predictive value (NPV) for detection of known resistance determinants, and their 95% confidence intervals (CI) were calculated with the epi.tests function of the epiR v1.0-14 package (53). False negative (FN) and false positive (FP) results were further investigated with alternative methods by querying the genome assemblies with Resfinder (54) and/or by mapping and local assembly of the sequence reads to the Bacterial Antimicrobial Resistance Reference Gene Database (Bioproject PRJNA313047) with ARIBA (55).

Seven studies reported ciprofloxacin MICs for a total of 889 *S.* Typhi strains, albeit interpreted with different breakpoint guidelines and versions (Additional File 2: Supplementary Table S1). We compared the Typhi Pathogenwatch ciprofloxacin resistance predictions (SIR) for each observed combination of genetic AMR determinants against the MIC values re-interpreted with the ciprofloxacin breakpoints for *Salmonella* spp. from CLSI M100 30^th^ edition (susceptible MIC≤0.06; intermediate MIC = 0.12 to 0.5; resistant MIC ≥1 (56)) with a script that is available at (49).

## Results

### Overview of the Typhi Pathogenwatch application

We have developed a public health focused application for *S.* Typhi genomics that uses genome assemblies to perform three essential tasks for surveillance and epidemiological investigations, i.e., (i) placing isolates into lineages or clonal groups, (ii) identifying their closest relatives and linking to their geographic distribution, and (iii) detecting the presence of genes and mutations associated with AMR. These data can aid the local investigator to rapidly identify a potential source of transmission and to predict AMR phenotypes.

The Pathogenwatch application can be accessed at https://pathogen.watch/styphi, where users can create an account and upload and analyse their genomes (Figure 1 and video (57)). User data remains private and stored in their personal account. Pathogenwatch provides compatibility with typing information for MLST (5), cgMLST (12), *in silico* serotyping (SISTR (15)), a SNP genotyping scheme (GenoTyphi (8)), and plasmid replicon sequences (36). The results for a single genome are displayed in a genome report that can be downloaded as a PDF. The results for a collection of genomes can be viewed online and downloaded as trees and tables of genotypes, AMR predictions, assembly metrics, and genetic variation. Results can also be accessed at a later date and shared via a collection ID embedded in a unique weblink, thus facilitating collaborative surveillance.

**Figure 1.**
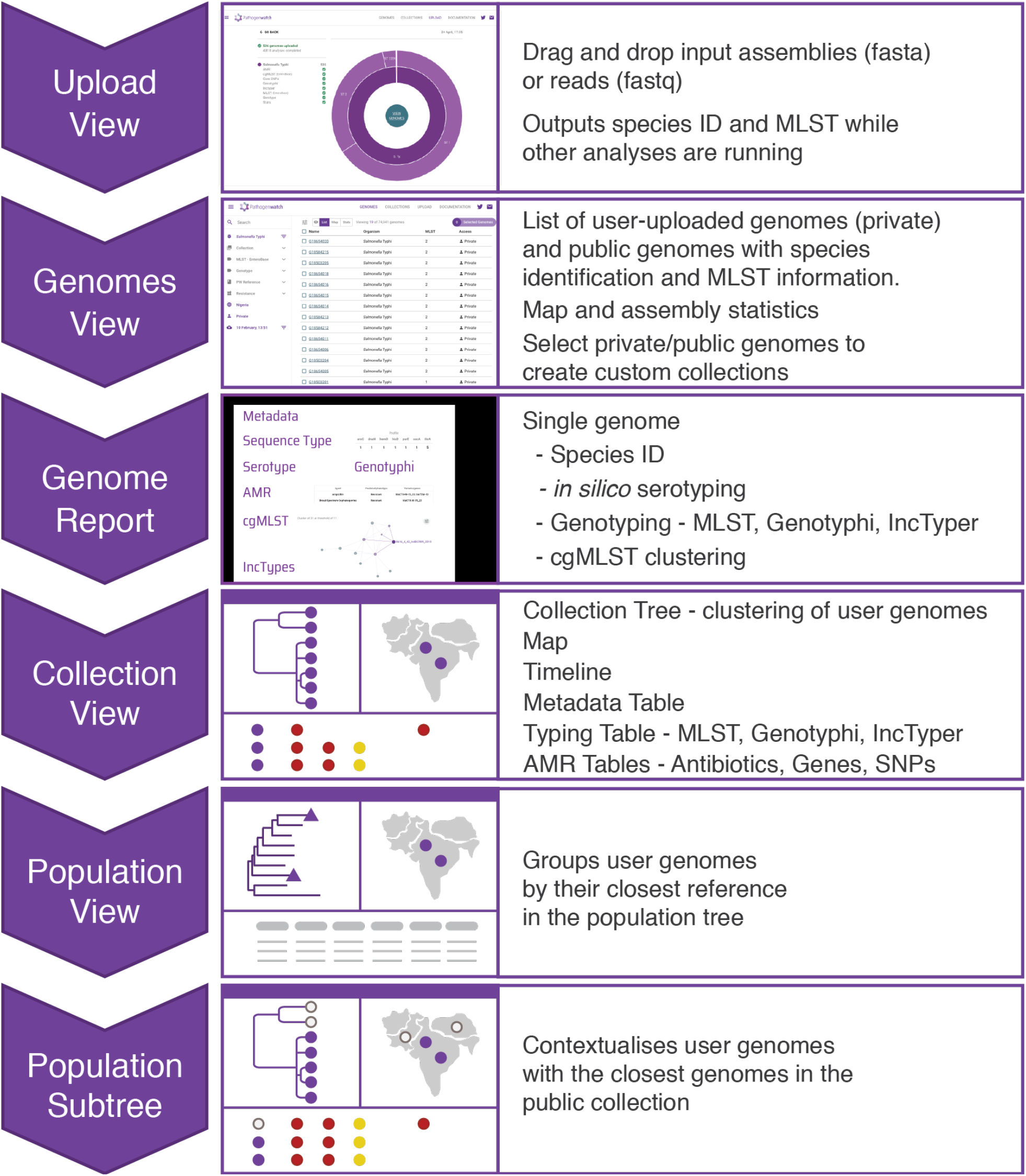
Workflow of the Typhi Pathogenwatch application. Input assemblies or sequence reads and metadata files can be uploaded via drag-and-drop onto the Upload page. Once the analyses completed, the genomes are listed on the Genomes page with Pathogenwatch outputs for speciation and MLST. Clicking on a genome name on the list pops up a Genome Report. The user can create collections of genomes. The Collection view displays the user genomes clustered by genetic similarity on a tree, their location on a map, a timeline, as well as tables for metadata, typing and AMR. The Population view displays the user genomes by their closest reference genome in the population tree. Clicking on one of the highlighted nodes (purple triangles) opens the Population subtree view, which contextualises the user genomes with the closest public genomes.

### Clustering genomes into lineages with Pathogenwatch

The pairwise genetic distance between isolates provides an operational unit for genomic surveillance. Typhi Pathogenwatch clusters genomes based on their genetic distance and displays their relationships in a collection tree. We benchmarked the Pathogenwatch clustering method against established methods of SNP-based tree inference, i.e. maximum likelihood or neighbour-joining trees inferred from core genome SNPs or whole-genome SNPs. We used three sets of published genomes: I) 118 genomes spanning the population diversity of *S.* Typhi defined by GenoTyphi (8); II) 138 closely related genomes, from a clonal expansion of 4.3.1 within Africa (7); and III) 43 strains from clade 3.2 including CT18, the genome of choice for reference-guided population genomics studies (8). The Pathogenwatch trees clustered the diverse genomes from subset I according to genotype assignments (Additional File 4: Supplementary Figure S1a), and detected phylogeographic signal in the closely related genomes of subset II (Additional File 4: Supplementary Figure S1b), in agreement with previous studies. In addition, we found that the Typhi Pathogenwatch clustering algorithm produced trees comparable to the established methods based on visualisations of the tree space and tree topology (Additional File 4: Supplementary Figure S2).

### Contextualisation with public data

A fundamental process for interpreting genomic datasets is to identify the nearest neighbours to the genome(s) under investigation. Pathogenwatch contextualises the user-uploaded genomes with public genomes using a population tree of 19 diverse genome references (Additional File 4: Supplementary Figure S3) to guide the SNP-based clustering of user and public genomes into subsets of closely related genomes (population subtrees). A previous investigation of a typhoid outbreak in Zambia exemplifies the value of contextualisation with the most relevant public data (58). This retrospective study identified clonal diversity and two repertoires of AMR genes within outbreak organisms, which belonged to haplotype H58 (genotype 4.3.1), but at the time only 5 genomes from 4.3.1 were available for comparison. Using Pathogenwatch, the outbreak strains can be rapidly contextualised with public genomes, which revealed two different clusters with close relationships to contemporary genomes from neighbouring countries Malawi and Tanzania (Figure 2a-b) that are also characterised by different *dfrA* genes (Figure 2c-d). The integration of genomic data and associated metadata from different studies in Pathogenwatch facilitates the investigation of a local outbreak in a broader geographic context via the web and without the need for bioinformatics expertise.

**Figure 2.**
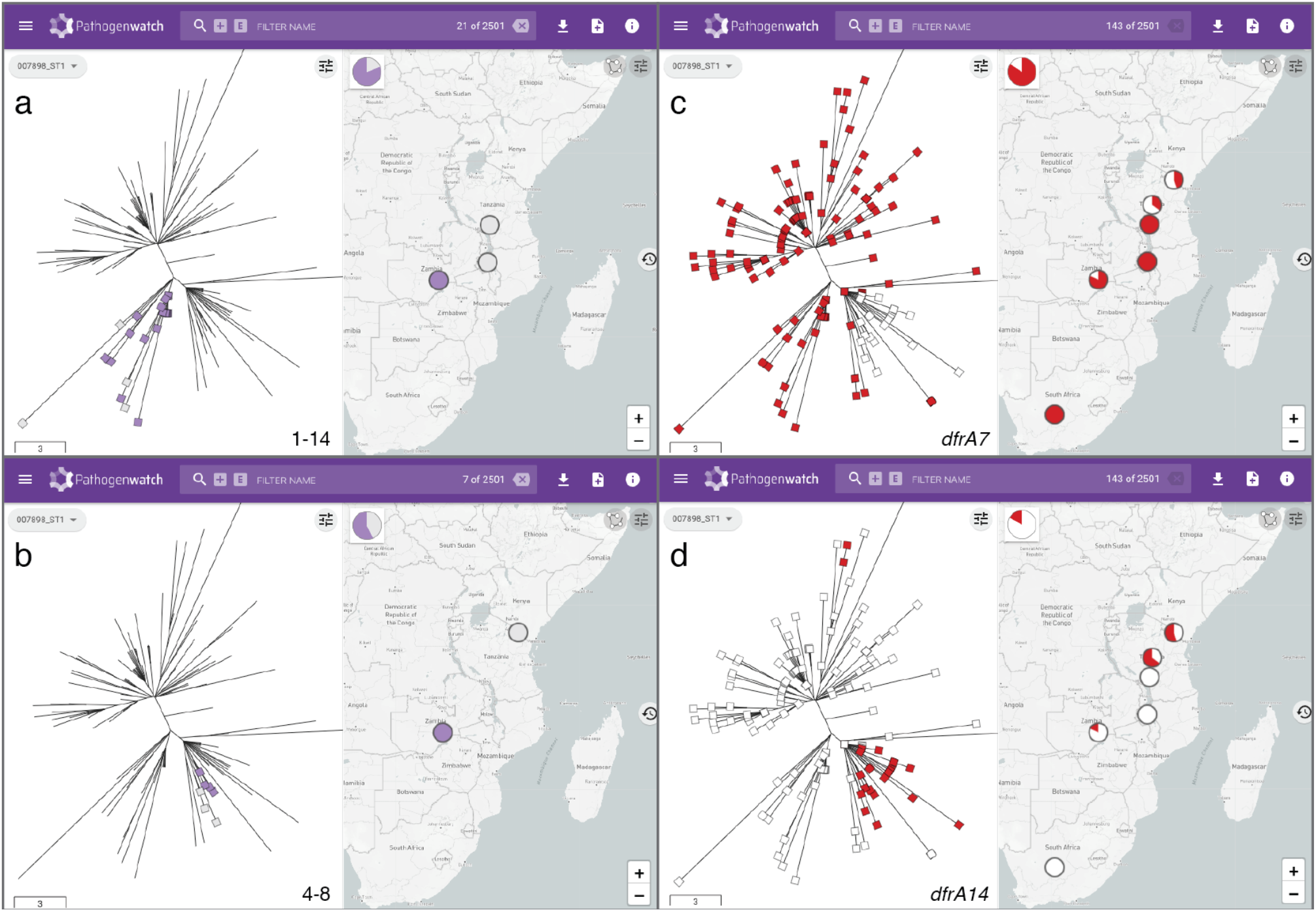
Pathogenwatch provides genomic context for outbreak investigations. **a-b** Genomes from an outbreak in Zambia (purple markers on tree and map) are linked by genetic relatedness to genomes from neighbouring countries Malawi and Tanzania (grey markers) forming 2 separate groups of 16 (**a**) and 4 (**b**) outbreak genomes, respectively. The number of pairwise differences (range) between outbreak and related genomes in the Pathogenwatch score matrix are indicated on the bottom-right of the tree panel. **c-d** Differential distribution of trimethoprim resistance genes *dfrA7* (**c**) and *dfrA14* (**d**) across the two clades containing outbreak genomes. The data are available at https://pathogen.watch/collection/g5pbucot6e58-hendriksen-et-al-2015.

Users interested in exploring the public genomes without creating their own collections can browse the public data as a whole (59) or view by published study (60). As of November 2020, Typhi Pathogenwatch included 4389 public genomes from 26 published articles (Additional File 1: Supplementary Table S1). The genomes spanned the years 1905 to 2019 and seventy-seven different countries, with the largest representation from 2000 onwards (N=3,795, 86.49%) and from the Indian subcontinent (N=1,602, 36.50%), respectively (Table 1 and Additional File 4: Supplementary Figure S4). Over 97% of the genomes were classified as either ST1 (68.2%) or ST2 (29.0%) using the 7-locus *Salmonella* MLST scheme, with the remaining 2.8% divided between 33 rare STs (Additional File 1: Supplementary Table S8). Similarly, over half of the genomes (N=2,500, 57.0%) belonged to the globally dominant MDR genotype 4.3.1, although the five different genotypes comprising 4.3.1 showed different temporal distributions and relative abundance (Additional File 4: Supplementary Figure S5).

**Table 1.** Characteristics of 4,389 public genomes in Pathogenwatch

| Year of isolation |  | Number of genomes (%) |
| --- | --- | --- |
|  | 1905-1969 | 41 (0.9) |
|  | 1970-1989 | 79 (1.8) |
|  | 1990-1999 | 395 (9.0) |
|  | 2000-2009 | 1,187 (27.0) |
|  | 2010-2019 | 2,609 (59.4) |
|  | No date | 78 (1.78) |
| Country of isolation (top 6) |  | Number of genomes (%) |
|  | Bangladesh | 637 (14.51) |
|  | United Kingdom | 629 (14.33) |
|  | India | 486 (11.07) |
|  | Nepal | 318 (7.25) |
|  | Vietnam | 220 (5.01) |
|  | Cambodia | 209 (4.76) |
| Assembly Stats |  | Median (range) |
|  | Number of contigs | 51 (1 – 633) |
|  | Assembly length | 4,747,975 (4,535,494 – 5,211,763) |
|  | N50 | 204,317 (19,527 – 4,806,333) |
|  | non-ATCG | 152 (0 – 48,002) |
|  | GC content (%) | 52.0 (51.4 - 52.4) |

### Genomic predictions of antimicrobial resistance

Typhi Pathogenwatch provides resistance predictions for antimicrobials relevant to treatment of typhoid fever by querying genome assemblies with BLAST (41) and a curated library of known AMR genes and mutations (Additional File 1: Supplementary Table S7). To benchmark the Typhi Pathogenwatch predictions, we compared the genotypic resistance genotypes to the available drug susceptibility phenotypes (SIR interpretation) of 1316 genomes, grouping the Resistant and Intermediate classifications as insusceptible. The sensitivity of the Pathogenwatch genotypic predictions was at least 0.96 for all antibiotics with a computed value (Table 2); at the time of writing, there were no insusceptible isolates described for colistin or meropenem. The false negative (FN) calls for ampicillin (N=4), cephalosporins (N=2), chloramphenicol (N=6), and sulfamethoxazole-trimethoprim (N=7) were paralled by the original genome studies (61–63), and by an alternative bioinformatics method (55), neither of which detected any known resistance genes in these genomes. The 49 FN calls for ciprofloxacin were also in agreement with the *in silico* analyses reported in the original genome studies (34, 63), in which no QRDR mutations or *qnr* genes were detected. Only mutations outside of the quinolone-resistance determining region (QRDR) of *parE* (A364V, N=17) or *gyrA* (D538N, N=2) were found in 20 genomes. These mutations have not as yet been shown to cause ciprofloxacin insusceptibility and were therefore omitted from the Pathogenwatch AMR library.

**Table 2.** Benchmark analysis of Typhi Pathogenwatch AMR predictions for ampicillin (AMP), chloramphenicol (CHL), broad-spectrum cephalosporins (CEP), ciprofloxacin (CIP), sulfamethoxazole-trimethoprim (SXT), tetracycline (TCY), azithromycin (AZM), colistin (CST) and meropenem (MEM). The total number of comparisons, true negatives (TN), true positives (TN), false negatives (FN), false positives (FN), sensitivity, specificity, positive predictive value (PPV), negative predictive value (NPV), very major error (VME) rate, major error (ME) rate, and concordance are shown. Confidence intervals (95%) are shown in parenthesis.

| Antibiotic | Total | TN | TP | FN | FP | Sensitivity<br>(95% CI) | Specificity<br>(95% CI) | PPV<br>(95% CI) | NPV<br>(95% CI) | VME<br>rate | ME<br>rate | Concordance<br>(%) |
| --- | --- | --- | --- | --- | --- | --- | --- | --- | --- | --- | --- | --- |
| AMP | 875 | 461 | 402 | 4 | 8 | 0.99<br>(0.97-1.00) | 0.98<br>(0.97-0.99) | 0.98<br>(0.96-0.99) | 0.99<br>(0.98-1) | 0.01 | 0.02 | 98.63 |
| CEP | 348 | 256 | 90 | 2 | 0 | 0.98<br>(0.92-1.00) | 1.00<br>(0.99-1.00) | 1.00<br>(0.96-1.00) | 0.99<br>(0.97-1.00) | 0.02 | 0 | 99.43 |
| CHL | 913 | 518 | 375 | 6 | 14 | 0.98<br>(0.97-0.99) | 0.97<br>(0.96-0.99) | 0.96<br>(0.94-0.98) | 0.99<br>(0.98-1.00) | 0.02 | 0.03 | 97.81 |
| CIP | 1282 | 111 | 1065 | 49 | 57 | 0.96<br>(0.94-0.97) | 0.66<br>(0.58-0.73) | 0.95<br>(0.93-0.96) | 0.69<br>(0.62-0.76) | 0.04 | 0.32 | 91.73 |
| SXT | 912 | 513 | 367 | 7 | 25 | 0.98<br>(0.96-0.99) | 0.95<br>(0.93-0.97) | 0.94<br>(0.91-0.96) | 0.99<br>(0.97- 0.99) | 0.02 | 0.05 | 96.49 |
| TCY | 44 | 40 | 4 | 0 | 0 | 1.00<br>(0.40-1.00) | 1.00<br>(0.91-1.00) | 1.00<br>(0.40-1.00) | 1.00<br>(0.91-1.00) | 0 | 0 | 100 |
| AZM | 156 | 144 | 12 | 0 | 0 | 1.00<br>(0.74-1.00) | 1.00<br>(0.97-1.00) | 1.00<br>(0.74-1.00) | 1.00<br>(0.97-1.00) | 0 | 0 | 100 |
| CST | 41 | 41 | 0 | 0 | 0 | - | 1.00<br>(0.91-1.00) | - | 1.00<br>(0.91-1.00) | - | 0 | 100 |
| MEM | 132 | 132 | 0 | 0 | 0 | - | 1.00<br>(0.97- 1.00) | - | 1.00<br>(0.97- 1.00) | - | 0 | 100 |

The specificity of the Pathogenwatch genotypic predictions was at least 0.95 for most antimicrobials (Table 2), with the exception of ciprofloxacin, for which a third of the ciprofloxacin susceptible isolates were reported as insusceptible by Pathogenwatch. A closer inspection of the 57 false positive (FP) results showed that Pathogenwatch reported one (N=55), two (N=2) or three (N=1) mutations in the QRDR of *gyrA, gyrB* and/or *parC*, most frequently the single mutations *gyrA*_S83F (N=25) and *gyrB*_S464F (N=16). For 54 of these samples, the same mutations were reported in the original genome studies. For the remaining three genomes, no mutations were reported in the original studies, but we confirmed the presence of *gyrB*_S464F (N=2) or *gyrB*_S464Y (N=1) in the assemblies using Resfinder (54). Similarly, we confirmed the Pathogenwatch identification of *bla*TEM-1, *catA1*, or *sul1*-*dfrA7* for all 47 of the FP calls for ampicillin (N=8), chloramphenicol (N=14), and sulfamethoxazole-trimethoprim (N=25), respectively, either from the original genome studies or with Resfinder.

The additive effect of QRDR mutations on ciprofloxacin susceptibility has been previously described (64). In addition, the presence of three non-synonymous mutations in the *gyrA* (S83F and D87N) and *parC* (S80I) genes was previously associated with ciprofloxacin resistance and fluoroquinolone treatment failure (64, 65) and was predictive of ciprofloxacin resistance in a study of reference laboratory isolates (66). Pathogenwatch thus reports this specific combination of mutations as resistant on the Antibiotics table with a red circle, while any other single, double or triple QRDR mutation is reported as decreased susceptibility (intermediate, yellow circle). We evaluated the ciprofloxacin MICs of 889 *S.* Typhi isolates from nine previous studies against the different combinations of resistance mechanisms identified by Pathogenwatch. Overall, the distribution of MIC values was consistent with the genomic predictions of AMR from Pathogenwatch (Figure 3). The isolates with 1 or 2 QRDR mutations displayed mostly intermediate MICs against ciprofloxacin, and support reporting as intermediate in Pathogenwatch. The MIC values of 7 isolates carrying single mutations on *gyrA* (S83F, S83Y) and *gyrB* (S464F), however, were below the intermediate breakpoint, consistent with the lower specificity reported for ciprofloxacin in Table 2. The highest ciprofloxacin MIC values were observed for the combination of *gyrA*_S83F-*gyrA*_D87N-*parC*_S80I mutations, reported as resistant by Pathogenwatch. However, the triple combination *gyrA*_S83F-*gyrA*_D87G-*parC*_E84K was represented by 9 isolates with MICs in both the resistant (N=6) and the intermediate (N=3) ranges, and is reported by Pathogenwatch as intermediate. Further susceptibility testing of isolates with this combination of mutations is needed to refine genotypic predictions. Likewise, several other mechanisms potentially conferring insusceptibility to ciprofloxacin were found in the public genomes but had with no or little associated MIC data, including seven additional triple mutations (Additional File 1: Supplementary Table S9, Additional File 4: Supplementary Figure S6).

**Figure 3.**
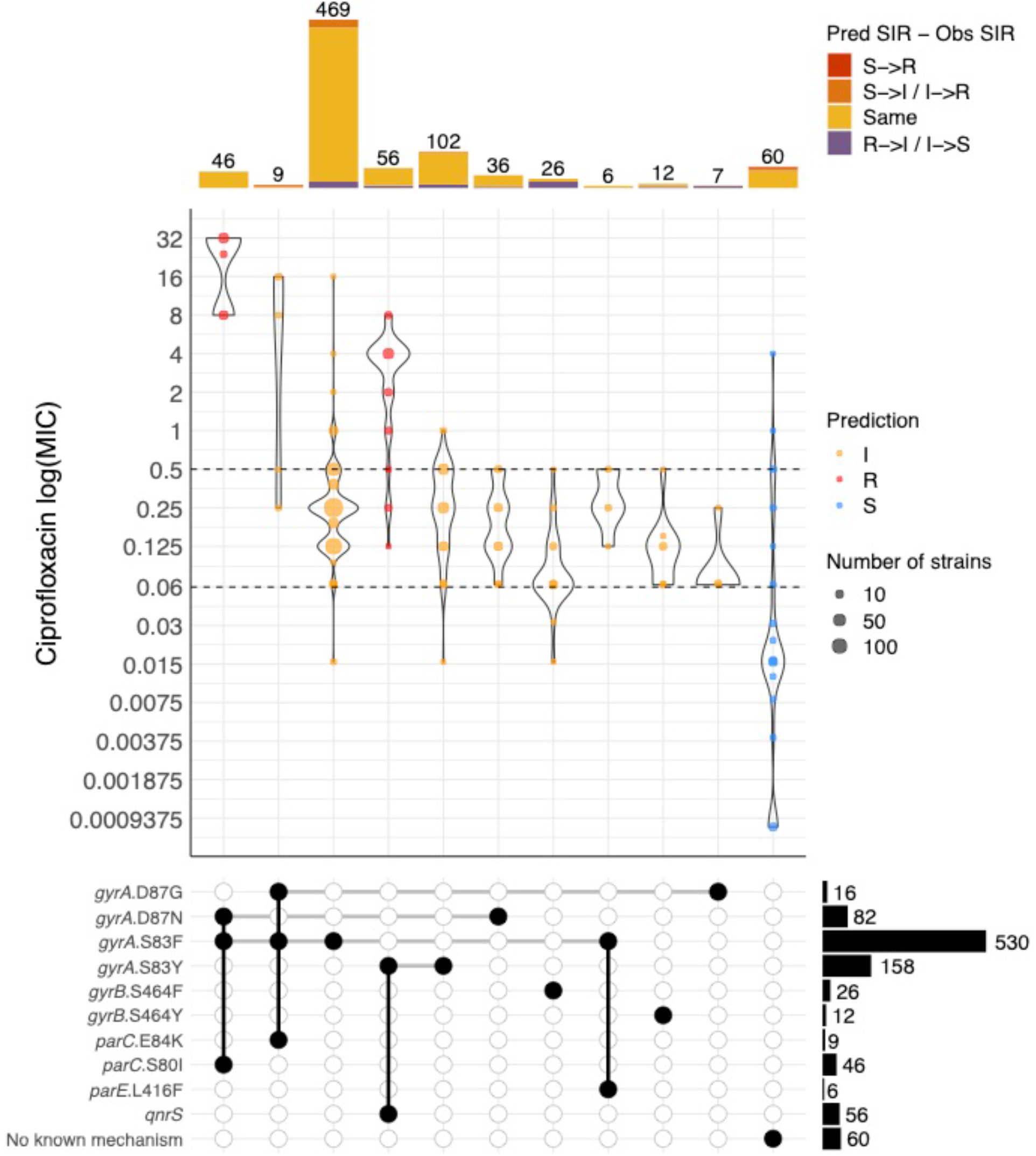
Distribution of minimum inhibitory concentration (MIC) values (mg/L) for ciprofloxacin in a collection of *S.* Typhi isolates with different combinations of genetic mechanisms that are known to confer resistance to this antibiotic. Only combinations observed in at least 5 genomes are shown. Dashed horizontal lines on the violin plots mark the CLSI clinical breakpoint for ciprofloxacin. Point colours inside violins represent the genotypic AMR prediction by Pathogenwatch on each combination of mechanisms. Barplots on the top show the abundance of genomes with each combination of mechanisms. Bar colours represent the differences between the predicted and the observed SIR (i.e. red for a predicted susceptible mechanism when the observed phenotype is resistant).

Genomic predictions of AMR are presented in three interactive and downloadable tables, Antibiotics, Genes, and SNPs, which display the predicted resistance profile, AMR genes and AMR-associated chromosomal SNPs found for each genome in the collection, respectively. The user can overlay the genotypic predictions on the tree and the map views for one or multiple antibiotics/genes/SNPs, thus intuitively linking resistance with genome clustering and geographic location. For example, the distribution of genomic predictions of ciprofloxacin resistant, MDR, or extremely drug resistant (XDR, defined as MDR + ciprofloxacin resistant) *S.* Typhi on the map and on the tree of 4389 public genomes highlight the lineages that represent a particular challenge to treatment and their geographical distribution (Additional File 4: Supplementary Figure S7). A summary of the genomic predictions of MDR and XDR *S.* Typhi highlights the differences in the distribution of high-risk clones by region, year and genotype, as inferred from the public genomes (Additional File 4: Supplementary Figure S8).

In addition, Pathogenwatch presents a granular picture of the different resistance mechanisms to an antibiotic. For example, the distinct distribution of trimethoprim-resistance gene *dfrA15* in West Africa associated with genotype 3.1.1, and of *dfrA7* across Central and East Africa, associated with genotypes 2.5.1 and 4.3.1.1, respectively (67) (Additional File 4: Supplementary Figure S9). The most frequent AMR genes in the public collection of 4389 genomes associated with an MDR phenotype were *bla*TEM-1 (ampicillin, N=1460), *catA1* (chloramphenicol, N=1406), *sul1* (sulfamethoxazole, N=1447), and *dfrA7* (trimethoprim, N=1232). Notably, *bla*CTX-M-15 was the most frequent gene coding for an extended-spectrum beta-lactamase (N=92, Additional File 4: Supplementary Figure S10). The acquired AMR genes found in the public genomes were identical or nearly identical matches to the AMR library representatives (Additional File 4: Supplementary Figure S11), with the vast majority of the matches (7842/8098, 96.8%) showing 100% identity.

Several plasmids have been implicated in the dissemination of drug-resistant *S.* Typhi. Notably, the MDR phenotype is linked to a composite transposon carrying multiple resistance genes, either located in IncH1 plasmids or integrated into the chromosome (7). An IncY plasmid that confers resistance to fluoroquinolones and third-generation cephalosporins was detected in XDR *S.* Typhi from an outbreak in Pakistan (61), while plasmids belonging to at least five different Inc types have been described in a recent pan-African study (67). Pathogenwatch identifies plasmid replicon marker sequences in the user genomes and reports them on the genome report and on the Typing table in the collection view (Figure 1). Pathogenwatch reported between one and four plasmid replicon marker sequences in a third of the public genomes (1,571/4,389, 35.79%, Additional File 4: Supplementary Figure S12a). Predictably, plasmid replicon markers were more frequent in genomes with predicted genotypic resistance, in particular those organisms that were resistant to multiple antimicrobials (Additional File 4: Supplementary Figure S12b). Notably, the cryptic plasmid pHCM2, which does not carry resistance genes (68), was the most common replicon detected amongst genomes in which acquired resistance genes were not detected. The distribution of replicon genes showed that the combination of IncH1A and IncH1B(R27) was prevalent in MDR genomes from Southeast Asia and East Africa belonging to clade 4.3.1, while the same combination with the addition of IncFIA(HI1) was more prevalent in West Africa, and associated with clade 3.1 (Additional File 4: Supplementary Figure 12b-d). The IncH1A and IncH1B(R27) sequences detect fragments of the *repA2* and *repA* genes, respectively, of the IncHI1 conjugative plasmid which has historically been associated with the majority of MDR typhoid (7). IncFIA(HI1) detects fragments of the *repE* gene that is present in a subset of IncHI1 plasmids, including the plasmid sequence type PST2 variant common in *S*. Typhi 3.1 in West Africa, but lacking from the PST6 variant that is widespread in *S*. Typhi 4.3.1 in East Africa and Asia (67).

### Maximising the utility of genomic data

Pathogenwatch makes the public WGS data easily accessible and searchable, and also constitutes a growing resource to which new data can be added. While genomic predictions of AMR are based on known mechanisms, the predictions can easily be updated as new mechanisms are discovered. Azithromycin is one of the last oral treatment options for typhoid for which resistance is currently uncommon, of particular importance in endemic areas with high rates of fluoroquinolone-resistance and outbreaks of XDR *S.* Typhi. A non-synonymous point mutation in the gene encoding the efflux pump AcrB (R717Q) was recently singled out as a molecular mechanism of resistance to azithromycin in *S*. Typhi (69). Pathogenwatch detected the *acrB*_R717Q mutation in a collection of 12 Bangladeshi genomes of genotype 4.3.1.1 isolated between 2013 and 2016 in which this mutation was first described (Figure 4). Notably, Pathogenwatch also detected the *acrB*_R717Q mutation in three additional genomes, two from isolates recovered in England in 2014 (no travel history available (70)), and one from an isolate recovered in Samoa in 2007 (7). The Samoan genome 10349_1_30_Sam072830_2007 was typed as genotype 3.5.4, while the English genomes 65343 and 32480 (no travel information available) belonged to genotypes 4.3.1.1 and 4.3.2.1, respectively. Genome 65343 was closely related to the cluster of 12 genomes from Bangladesh where this mutation was first described, while genome 32480 belonged to a small cluster of five genomes from India or with travel history to India. Thus, reanalysis of public data with Pathogenwatch showed that the *acrB*_R717Q mutation has emerged in multiple genetic backgrounds, in multiple locations, and as early as 2007.

**Figure 4.**
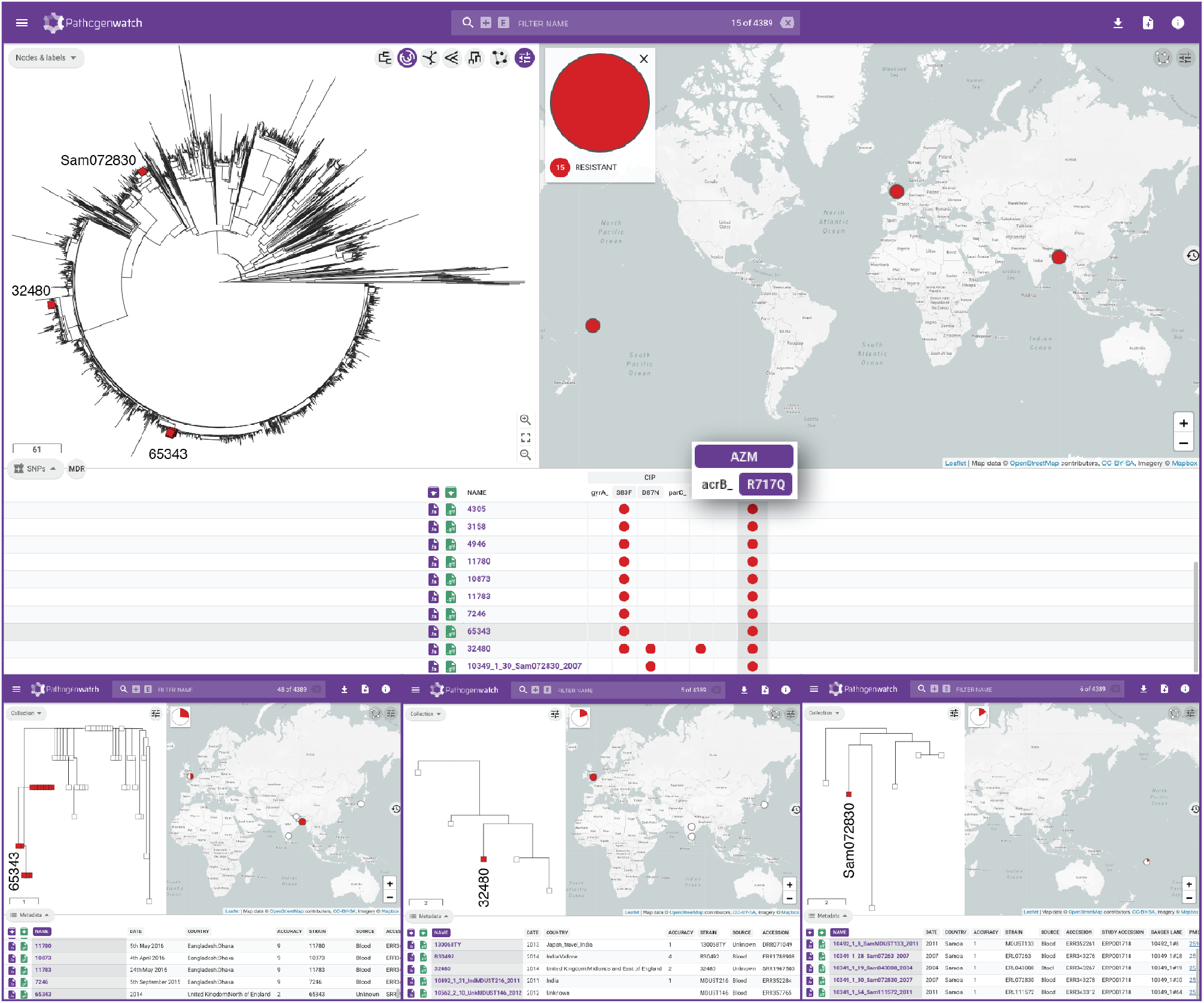
Pathogenwatch data reusability. Fifteen genomes carrying the *acrB*_R717Q mutation recently linked to azithromycin resistance in *S.* Typhi are shown in red on the tree of 4389 public genomes and on the map. The presence of the mutation is indicated by the red circles on the SNPs table. Three of these genomes (tree labels) belong to isolates collected before the mutation was first described and are shown in more detail in the bottom panels. The data are available at https://pathogen.watch/collection/07lsscrbhu2x-public-genomes

### Pathogenwatch applied to rapid risk assessment

Typhoid fever is rare in countries with a good infrastructure for the provision of clean water and sanitation, with most cases arising from travel to endemic areas (71). Ceftriaxone-resistant typhoid fever was recently reported in developed countries and associated with travel to Pakistan (72–74). These ceftriaxone resistant isolates were associated to the recent outbreak of XDR *S.* Typhi in the Sindh province of Pakistan by the epidemiological data, the antibiograms, and information derived from WGS of the clinical isolate, such as presence of resistance genes, and mobile genetic elements. In some cases the genomes were contextualised with retrospective genomes by building a phylogenetic tree using an existing bioinformatic pipeline (72, 73).

We exemplify how Pathogenwatch facilitates this analysis with the genome from the isolate recovered in Canada (PHL5950, accession RHPM00000000 (74)). Pathogenwatch provides a printable genome report (Additional File 4: Supplementary Figure S13) including genotyping and *in silico* serotyping information, predicted resistance profile, and the presence of resistance genes and mutations. In addition, Pathogenwatch places the genome within the Pakistani XDR outbreak (Figure 5) and shows the close genetic relatedness (between 3 and 8 pairwise differences) of the isolates via the downloadable score matrix.

**Figure 5.**
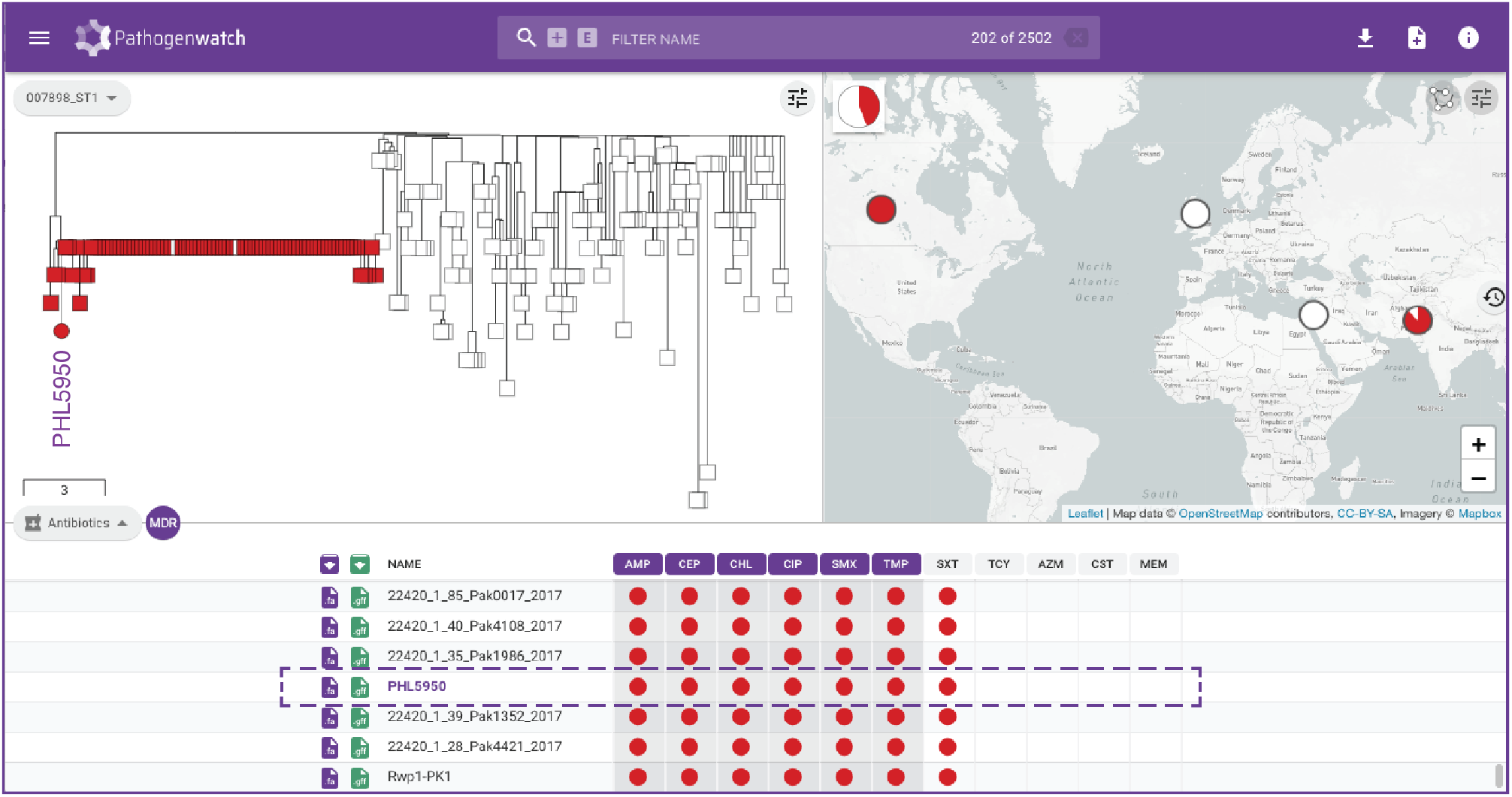
Rapid risk assessment of typhoid fever cases in non-endemic regions. Pathogenwatch places genome PHL5950 from an isolate recovered in Canada and with travel history to Pakistan within the XDR-outbreak in Pakistan (red markers). The data are available at https://pathogen.watch/collection/11lsok8nrzts-wong-et-al-2018-idcases-15e00492

### Pathogenwatch as a tool for international collaboration in typhoid surveillance

As WGS capacity becomes a reality in typhoid endemic countries, there is a growing opportunity for local genomic surveillance and for collaboration across borders. This is underscored by the growing number of genomes from the Indian Subcontinent (Additional File 4: Supplementary Figure S3), where epidemic clone 4.3.1 (H58) and the nested clade of fluoroquinolone-resistant triple mutants belonging to genotype 4.3.1.2 (H58 lineage II) have been shown to have originated (7, 65)). The triple mutants were first reported in Nepal (isolated in 2013-2014) and linked to isolates from India from 2008-2012 (65). More recent surveillance studies showed that this lineage was still prevalent in *S.* Typhi isolates collected in Nepal in 2016 and in India in 2016-2017 (34, 75). The public data integrated in Pathogenwatch showed that (at the time of writing) this lineage is represented by 195 public genomes from seven countries (India, Bangladesh, Nepal, Pakistan, Myanmar, Japan, and United Kingdom, Figure 6a, (7, 63, 64, 70, 75–78)) and from as early as 2006 (Japan, with travel history to India, Figure 6b (76)). Linking the tree and the map highlights distinct clusters of genomes that show evidence of transmission across borders, for example between India-Pakistan and India-Nepal (Figure 8c-d). In addition, three isolates recovered in 2016 in India were reported to be resistant to cephalosporins, linked to the presence of the *bla*_SHV-12_ gene (79); Pathogenwatch detected *bla*_SHV-12_, *qnrB* and the IncX3 plasmid replicon in these genomes. Another previous study reported an IncN replicon in three genomes from the United Kingdom (two with travel history to India) that also carried resistance genes *dfrA15* (trimethoprim), *sul1* (sulfamethoxazole), and *tetA*(A) (tetracycline) (64). Pathogenwatch identified the same AMR genes and plasmid replicon in these genomes, and also in two closely related genomes from Japan with travel history to Nepal and India (Figure 6b). Altogether, these observations suggest that this lineage circulating in South Asia and linked to treatment failure with fluoroquinolones, can acquire plasmids with additional AMR genes, with the concomitant risk of the clonal expansion of a lineage that poses additional challenges to treatment.

**Figure 6.**
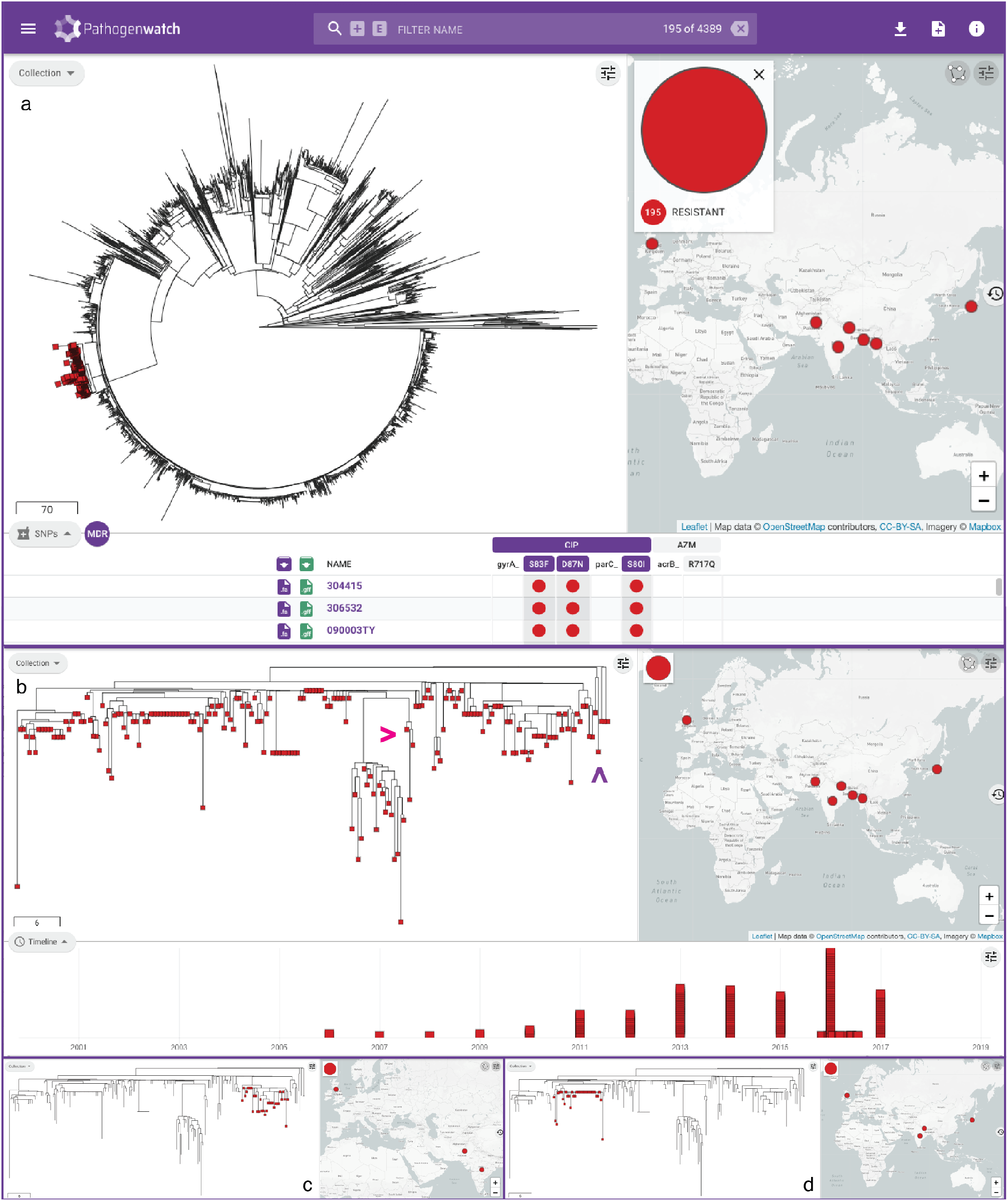
Pathogenwatch to for collaborative international surveillance of *S.* Typhi. **a** Pathogenwatch highlights 195 ciprofloxacin-resistant triple mutants on the public data tree and map by simultaneously selecting the mutations *gyrA*_S83F, *gyrA*_D87N, and *parC*_S80I on the SNPs table**. b** Detailed visualisation of the triple mutants showing the temporal distribution of the genomes on the timeline. Magenta arrowhead: 3 genomes from India with *bla*_SHV-12_, *qnrB* and an IncX3 replicon. Purple arrowhead: 4 genomes with *sul1*, *dfrA15*, *tetA*(A) and an IncN replicon from the UK and Japan. Selecting individual clades on the tree shows distinct clades that span neighbouring countries India-Pakistan (**c**) and India-Nepal (**d**). The data are available at https://pathogen.watch/collection/07lsscrbhu2x-public-genomes

## Discussion

Our understanding of the *S.* Typhi population structure, including MDR typhoid has improved dramatically since the introduction of WGS, which provides a level of discrimination much needed for a human-adapted pathogen that exhibits very limited genetic variability. Progress towards the widespread implementation of WGS for epidemiological investigations and integrated routine surveillance within public health settings needs to be accompanied by i) active surveillance programs in endemic regions; ii) implementation of WGS at laboratories in endemic regions; iii) analysis of WGS data with fast, robust and scalable tools that deliver information for public health action; iv) dissemination of WGS data through networks of collaborating reference laboratories at the national, international and global scales; and v) provision of WGS data and associated metadata through continuously growing databases that are amenable to interaction and interpretation (80). Here, we described Typhi Pathogenwatch, a web application for the genomic surveillance and epidemiology of *S.* Typhi, which enhances the utility of public WGS data and associated metadata by integration into an interactive resource that users can browse or query with their own WGS data.

Rapid, timely access to information on local patterns of AMR may inform treatment regimens, which could ultimately lead to a reduction in morbidity and mortality associated with enteric fever as this is much greater in the absence of effective antimicrobial therapy (81). Typhi Pathogenwatch provides a general framework for genomic predictions of AMR and of related strain clusters, and is accessible to users of all bioinformatics skills levels. This enables users with an understanding of genomics but no bioinformatics training to conduct surveillance and epidemiological investigations using WGS. Furthermore, it allows experienced bioinformaticians to rapidly perform the essential tasks listed in the results section, thus freeing up time for more advanced downstream analyses.

We demonstrated that genomic predictions of AMR are largely concordant with the resistance phenotype (overall concordance 96.34%, Table 2). It should also be noted that Pathogenwatch was developed with a focus on surveillance, not for clinical decision making. A previous study of 332 *S.* Typhi isolates analysed in a single reference laboratory reported only 0.03% discordant results (66) versus 3.66% from our data. Similarly, AMRFinder (11) and Resfinder 4.0 (10) reported slightly higher overall phenotype-genotype concordance, at 98.0% and 98,8%, respectively. These two studies analysed large collections of *Salmonella* genomes, albeit belonging to non-typhoidal serovars. A limitation of our study, is that it amalgamated published susceptibility data from thirteen different publications conducted in eight different countries. The availability of consistent laboratory antimicrobial susceptibility testing data is key for the periodic benchmarking and refinement of genomic predictions of AMR (82), in particular for ciprofloxacin due to the diverse combinations of mechanisms (Additional File 1: Supplementary Table S9). Unlike other AMR prediction tools, Pathogenwatch provides the added value of immediate contextualisation with location, time and population structure in an interactive visualisation.

Novel mechanisms of AMR can easily be added to the curated Pathogenwatch AMR library, and the growing collection of public genomes can be retrospectively screened, potentially revealing the presence of a newly identified gene or mutation in genomes from isolates previously collected (Figure 4). This illustrates how the provision of public genomic data through Pathogenwatch maximises reusability from which new insights into novel AMR mechanisms can be derived. The utility of maintaining a regularly updated archive of WGS data that can be rapidly ‘mined’ for the presence of newly discovered AMR gene was elegantly illustrated before by the retrospective discovery of the colistin resistance gene *mcr-1* in *S. enterica* and *Escherichia coli* genomes from Public Health England (83). Pathogenwatch extends this utility to the entire Typhi community, thus democratising the reusability of the genomic data.

Contextualizing new genomes with existing data is now a routine part of genomic epidemiology, as it can complement epidemiological investigations to, among many applications, place the new genomes in or out of an outbreak, link to past outbreaks, and determine if the success of a resistant phenotype is the result of a single clonal expansion or multiple independent introductions (84). Using the publicly available genomes, we provided examples of the utility of Pathogenwatch to contextualise user-uploaded genomes for outbreak investigation in endemic areas (Figure 2) or for the management of patients in non-endemic countries with travel history to endemic areas (Figure 5). Analysing new genomes in the context of global genomes involves the retrieval, storage and bioinformatic analysis of large amounts of sequence data and linked metadata, which is time-consuming at the least, and largely unfeasible for hospitals or public-health agencies with limited computing infrastructure. Pathogenwatch bridges this gap and provides contextualisation with the closest genomes guided by the *S.* Typhi population tree (Additional File 4: Supplementary Figure S3) and subtrees.

The interpretation of the genomic context relies heavily on the completeness of the public collection used for contextualisation and of its metadata. This in turn depends on the establishment of local, national and international surveillance programs for the real-time management of emerging lineages that pose a direct threat to human health. The International Typhoid Consortium collected and sequenced around 40% of the global genomes available in Pathogenwatch for comparison (7, 8), but ongoing surveillance and WGS are needed to maintain a relevant, contemporary genome collection. Pathogenwatch does not currently support automated updates or submissions, which instead requires retrieval and curation of published genome data and associated metadata. Thus, while sequence data are not instantly available on Pathogenwatch when they become available on sequence data archives. Pathogenwatch maximises the utility of genomes available on the platform. For example, as of November 2020 Pathogenwatch provides 4234 of 4389 (96.5%) *S.* Typhi genomes with at least both year and country of isolation, while the same applies to 3473 of 7743 (44.9%) genomes on Enterobase (16), 3936 of 5618 (70.1%) genomes on GenomeTrakr (14), and 2085 of 3100 (67.3%) genomes on PATRIC (13). In addition, Pathogenwatch includes patient travel information when available.

Pathogenwatch can facilitate collaborative surveillance in endemic areas via data integration and shared collections for the early detection and containment of high-risk clones (Figure 6). Collections can be set to off-line mode to work while disconnected from the internet, which may be advantageous in settings with unreliable internet connections. Despite recent efforts to promote data openness in times of pandemics (85, 86), several challenges to sharing genomic data and linked metadata remain in both the academic and public-health settings (80). User-uploaded genomes and metadata remain in the Pathogenwatch user account, and collections also remain private unless the user specifically shares them via a collection URL. Moreover, Pathogenwatch offers a private metadata option to work with confidential information.

Recent improvements in our understanding of the disease burden and the dissemination of AMR, and the development of new typhoid conjugate vaccines have bolstered efforts to employ routine vaccination for the containment of typhoid fever (87). Routine surveillance coupled with WGS can inform decisions on suitable settings for the introduction of vaccination programs and on the evolution of pathogens in response to them (88, 89). Pathogenwatch could be linked to routine genomic surveillance around typhoid vaccination initiatives to monitor the population dynamics in response to the deployment of new vaccines.

## Conclusions

Typhi Pathogenwatch combines accurate genomic predictions of AMR with broad geographic and population context within an easy to use interface for delivered for the community and to support ongoing typhoid surveillance programs. The modular architecture of Pathogenwatch allows new functionalities to be added to cater to the community needs.

## List of abbreviations

AMR: antimicrobial resistance
cgMLST: core-genome multi-locus sequence typing
PFGE: pulse-field gel electrophoresis
MDR: multi-drug resistant
MLST: multi-locus sequence typing
MLVA: multiple-locus variable-number tandem-repeat analysis
QRDR: quinolone resistance determining region
VNTR: multiple-locus variable-number tandem-repeat
XDR: extremely-drug resistant
WGS: whole-genome sequencing

## Availability of data and materials

The genome data and linked metadata presented are available from: https://pathogen.watch/collection/07lsscrbhu2x-public-genomes, https://pathogen.watch/collection/g5pbucot6e58-hendriksen-et-al-2015, and https://pathogen.watch/collection/11lsok8nrzts-wong-et-al-2018-idcases-15e00492

The tree comparison nexus files are available from https://gitlab.com/cgps/pathogenwatch/publications/styphi/benchmark_tree

The AMR benchmarking input files and script are available from https://gitlab.com/cgps/pathogenwatch/publications/styphi/benchmark_AMR

## Competing interests

The authors declare no competing interests.

## Funding

Pathogenwatch is developed with support from Li Ka Shing Foundation (Big Data Institute, University of Oxford) and Wellcome (grant number 099202). SA and DMA are supported by the National Institute for Health Research (UK) Global Health Research Unit on genomic Surveillance of AMR (16_136_111) and by the Centre for Genomic Pathogen Surveillance (http://pathogensurveillance.net).

ZAD received funding from the European Union’s Horizon 2020 research and innovation programme under the Marie Skłodowska-Curie grant agreement TyphiNET No 845681.

## Authors’ contributions

DMA conceived the Pathogenwatch application. CY, RJG, KA, BT, AU and DMA developed the Pathogenwatch application. SA drafted the manuscript. SA, DMA, KEH, SB, and GD contributed to the conception and design of the work, data interpretation, and substantially revised the manuscript. SA, CY, VKW, ZAD, SN, AJP, JAK, SEP and FM contributed to the acquisition and interpretation of data. SA, CY and LSB analysed the data. All authors read and approved the final manuscript.

## Acknowledgements

We are grateful to Flora Stevens and Joanne Freedman from the Travel Health and IHR department at Public Health England for providing some of the travel information linked to isolates from the United Kingdom, and to Dr. Koji Yahara, Dr. Makoto Ohnishi and Dr. Masatomo Morita for providing the travel information linked to isolates from Japan.

## Notes

https://gitlab.com/cgps/pathogenwatch/publications/styphi/

### Competing Interest Statement

The authors have declared no competing interest.

