## Additional File 1 Supplementary Tables for "A global resource for genomic predictions of antimicrobial resistance and surveillance of *Salmonella* Typhi at Pathogenwatch"

**Supplementary Table S1.** Collections of *S.* Typhi genomes included in Pathogenwatch at the time of writing. The availability of strain-level antimicrobial susceptibility testing (AST) data and of MIC values for ciprofloxacin is indicated.

| Collection name | PMID | No. of Genomes | Location | AST performed/  data used for concordance | CIP MIC data available | Reference |
| --- | --- | --- | --- | --- | --- | --- |
| Ahmad et al. (2017) | 28254988 | 1 | Malaysia | yes / no | no | (1) |
| Ashton et al. (2016) | 27069781 | 513 | UK (Travel multi) | no | no | (2) |
| Baker et al. (2015) | 26411565 | 37 | NA | no | no | (3) |
| Britto et al. (2018) | 29684021 | 192 | Nepal | yes / yes | no | (4) |
| Britto et al. (2020) | 31665304 | 94 | India | yes / yes | no | (5) |
| Burnsed et al. (2018) | 30236166 | 30 | USA  (Travel Marshall Islands) | yes / no | no | (6) |
| Djeghout et al. (2018) | 29616895 | 1 | Bangladesh | yes / yes | yes | (7) |
| Dyson et al. (2017) | 28060810 | 45 | Thailand | no | no | (8) |
| Gul et al. (2017) | 29051234 | 1 | Pakistan | yes / no | no | (9) |
| Hendriksen et al. (2015) | 25392358 | 22 | Zambia | yes / no | no | (10) |
| Hendriksen et al. (2015) | 25428145 | 2 | Netherlands and Norway (Travel Philippines) | yes / yes | yes | (11) |
| Hooda et al. (2019) | 31730615 | 12 | Bangladesh | yes / yes | no | (12) |
| Ingle et al. (2019) | 31513580 | 524 | UK (Travel multi) | yes (N=173) / yes | yes | (13) |
| Klemm et al. (2018) | 29463654 | 100 | Pakistan | yes / yes | no | (14) |
| Matono et al. (2017) | 29255729 | 93 | Japan (Travel multi) | yes / yes | no | (15) |
| Oo et al. (2019) | 31225619 | 39 | Myanmar | yes / yes | yes | (16) |
| Park et al. (2018) | 30504848 | 249 | Africa multi | no | no | (17) |
| Pham Thanh et al. (2016) | 26974227 | 78 | Nepal | yes / yes | yes | (18) |
| Phoba et al. (2017) | 29136410 | 1 | DR Congo | yes/no | no | (19) |
| Pragasam et al. (2020) | 32003431 | 194 | India | yes (N=30) / yes | no | (20) |
| Rodrigues et al. (2017) | 25961941 | 3 | India | yes / no | no | (21) |
| Sah et al. (2019) | 31872221 | 2 | India | yes / yes | yes | (22) |
| Tanmoy et al. (2018) | 32003431 | 536 | Bangladesh | yes / yes | yes | (23) |
| Wong et al. (2015) | 25961941 | 1822 | Multi | no | no | (24) |
| Wong et al. (2016) | 27703135 | 99 | Multi | no | no | (25) |
| Wong et al. (2016) | 27657909 | 128 | Nigeria | yes / no | no | (26) |

**Supplementary Table S3.** Reference genomes used to compute the Pathogenwatch *S.* Typhi core genome library (N=26) and the population tree (N=19).

| Strain Name | Country | Collection Year | MLST ST | GenoTyphi genotype | Assembly Length | N50 | No. contigs | Non-ATCG | % GC Content | % Core Families | Accessions | Study Accession | PMID | Core Library | Population Tree Reference |
| --- | --- | --- | --- | --- | --- | --- | --- | --- | --- | --- | --- | --- | --- | --- | --- |
| E02-1180 | India | 2002 | 2 | 0.0.3 | 4838914 | 4793071 | 4 | 0 | 52.1 | 100 | ERR581073,  ERR581085 | PRJEB5919 | 25961941 | yes | yes |
| E00-7866 | Morocco | 2000 | 2 | 0.1 | 4831122 | 4797408 | 3 | 0 | 52.1 | 100 | ERR581072,  ERR581087 | PRJEB5919 | 25961941 | yes | yes |
| 76-1292 | Democratic Republic of Congo | 1976 | 8 | 1.1.3 | 4934685 | 4780549 | 3 | 0 | 52.1 | 100 | ERR601541, ERR601549 | PRJEB5919 | 25961941 | yes | yes |
| M326 | Unknown | 1939 | 2 | 1.2.1 | 4775555 | 204340 | 47 | 8 | 52 | 100 | ERR279243 | PRJEB3255 | 26411565 | no | yes |
| 11909_3 | Mexico | 2011 | 2 | 2.0.2 | 4811824 | 4753305 | 4 | 0 | 52 | 100 | ERR584382,  ERR584388 | PRJEB5919 | 25961941 | yes | yes |
| E98-3139 | Mexico | 1998 | 2 | 2.0.2 | 4675865 | 3003486 | 6 | 0 | 52.1 | 100 | ERR581098, ERR581108, ERR601543 | PRJEB5919 | 25961941 | yes | no |
| ERL024120 | Indonesia | 2002 | 2138 | 2.1.2 | 4811886 | 4811886 | 1 | 0 | 52.1 | 100 | ERR581076, ERR581091 | PRJEB5919 | 25961941 | yes | yes |
| M223 | Unknown | 1939 | 2 | 2.1.7 | 4849011 | 4806333 | 3 | 0 | 52.1 | 100 | ERR581077, ERR581088 | PRJEB5919 | 25961941 | yes | no |
| 80-2002 | Madagascar | 1980 | 2 | 2.2 | 5061131 | 4819925 | 4 | 0 | 52.1 | 100 | ERR601542, ERR601550 | PRJEB5919 | 25961941 | yes | no |
| ERL114000 | Nepal | 2011 | 2 | 2.2 | 4776059 | 4776059 | 1 | 0 | 52.1 | 100 | ERR581075, ERR581086 | PRJEB5919 | 25961941 | yes | no |
| UI4692 | Laos | 2004 | 2 | 2.2.2,2.2.3 | 4707798 | 185395 | 43 | 0 | 52.1 | 100 | ERR331348 | PRJEB3215 | 25961941 | no | yes |
| ERL103914 | South America | 2010 | 2 | 2.3.2 | 4789623 | 4789623 | 1 | 0 | 52.1 | 100 | ERR581074, ERR581092 | PRJEB5919 | 25961941 | yes | no |
| ST1921-06 | Argentina | 2006 | 2 | 2.3.3 | 4718006 | 204238 | 42 | 0 | 52.1 | 100 | ERR353335 | PRJEB3215 | 25961941 | no | yes |
| LNT1360 | Laos | 2010 | 2218 | 2.4 | 4794285 | 206322 | 40 | 2 | 52.1 | 100 | ERR326664 | PRJEB3215 | 25961941 | yes | no |
| D50739 | Malawi | 2009 | 2 | 2.4.1 | 4704071 | 206289 | 45 | 24 | 52.1 | 100 | ERR279161 | PRJEB3215 | 25961941 | yes | yes |
| E98-0664 | Kenya | 1998 | 2 | 2.5 | 4764488 | 4752533 | 3 | 0 | 52.1 | 100 | ERR601554, ERR601551 | PRJEB5919 | 25961941 | yes | yes |
| H12ESR00755-001A | Philippines | 2012 | 1 | 3 | 4814227 | 4802146 | 2 | 0 | 52.1 | 100 | ERR581080, ERR581089 | PRJEB5919 | 25961941 | yes | yes |
| Quailes | USA | 1958 | 2 | 3.1 | 4841927 | 4797293 | 4 | 0 | 52.1 | 100 | ERR634098, ERR634100, ERR654508 | PRJEB5919 | 25961941 | yes | no |
| 404Ty | Indonesia | 1983 | 2 | 3.1.2 | 4890679 | 4807711 | 6 | 0 | 52 | 100 | ERR581078, ERR581090 | PRJEB5919 | 25961941 | yes | yes |
| CT18 | Vietnam | 1993 | 2 | 3.2.1 | 5133713 | 4809037 | 3 | 0 | 51.9 | 100 | GCA_000195995.1 | PRJNA236 | 11677608 | yes | yes |
| ERL041834 | India | 2004 | 2 | 3.3 | 4892436 | 4675147 | 5 | 0 | 52.1 | 100 | ERR581081, ERR581083 | PRJEB5919 | 25961941 | yes | yes |
| H12ESR04734-001A | India | 2012 | 2 | 3.3 | 4943859 | 4876386 | 5 | 0 | 52.1 | 100 | ERR581099, ERR581105 | PRJEB5919 | 25961941 | yes | no |
| 002168 | Cambodia | 2010 | 1 | 3.4 | 4764266 | 217780 | 42 | 2 | 52.1 | 100 | ERR360747 | PRJEB3215 | 25961941 | yes | no |
| LNT1148 | Laos | 2001 | 1 | 3.4 | 4743503 | 204320 | 46 | 2 | 52.1 | 100 | ERR340779 | PRJEB3215 | 25961941 | yes | yes |
| ERL024919 | Samoa | 2002 | 1 | 3.5.4 | 4805266 | 4784637 | 2 | 0 | 52.1 | 100 | ERR581094, ERR581102 | PRJEB5919 | 25961941 | yes | no |
| H12ESR00394-001A | Samoa | 2012 | 1 | 3.5.4 | 4808147 | 4808147 | 1 | 0 | 52.1 | 100 | ERR581095, ERR581103 | PRJEB5919 | 25961941 | yes | yes |
| Ty2 | Russia | 1916 | 1 | 4.1 | 4791961 | 4791961 | 1 | 11 | 52.1 | 100 | GCA_000007545.1 | PRJNA371 | 12644504 | yes | yes |
| ERL072973 | Fiji | 2007 | 1 | 4.2.2 | 4806043 | 4789877 | 2 | 0 | 52.1 | 100 | ERR581097, ERR581107 | PRJEB5919 | 25961941 | yes | yes |
| 007898 | Cambodia | 2010 | 1 | 4.3.1.1 | 4793925 | 4793925 | 1 | 0 | 52.1 | 100 | ERR752439, ERR752446 | PRJEB5919 | 25961941 | yes | yes |

**Supplementary Table S7.** List of genetic AMR determinants in Typhi Pathogenwatch. Effect: R = Resistance, I = Intermediate resistance (decreased susceptibility). Genes and mutations in bold type have been demonstrated to have a role on AMR in *S.* Typhi, the remaining mechanisms are of emerging importance in Gram negative bacteria. Pathogenwatch Library 1: *S.* Typhi AMR (27), 2: ESBL (28), 3: Carbapenemases (29), 4: Colistin resistance (30)

| Antibiotic class | Antibiotic (abbreviation used in Pathogenwatch) | Genetic determinants of resistance | Effect | Pathogenwatch Library (References) |
| --- | --- | --- | --- | --- |
| Phenicols | Chloramphenicol (CHL) | ***catA1*** | R | 1 (31) |
|  |  | ***cmlA*** | R | 1 (32) |
| Fluoroquinolones | Ciprofloxacin (CIP) | ***qnrS*, *qnrB***, *qnrA*, *qnrD* | I | 1 (33) |
|  |  | ***gyrA* S83F, S83Y, D87N, D87Y, D87A, D87G, D87V** | I | 1 (33, 34) |
|  |  | ***gyrB* S464F, S464Y, Q465R, Q465L** | I | 1 (33) |
|  |  | ***parC* S80I, E84G, E84K** | I | 1 (33) |
|  |  | ***parE* L416F, D420N** | I | 1 (33, 35, 36) |
| Sulfonamides | Sulfamethoxazole (SXM) | ***sul1*, *sul2*** | R | 1 (32) |
| Folate Pathway Inhibitors | Trimethoprim (TMP) | ***dfrA1*, *dfrA5*, *dfrA7*, *drfA14*, *dfrA15*, *dfrA17*, *dfrA18*** | R | 1 (32) |
|  | Tetracycline (TCY) | ***tetA*(A), *tetA*(B), *tetA*(C), *tetA*(D)** | R | 1 (31) |
| Macrolides | Azithromycin (AZM) | *ermA*, *ermB*, *ermC* | R | 1 |
|  |  | *ereA*, *ereB* | R | 1 |
|  |  | *mefA* | R | 1 |
|  |  | *mphA*, *mphB* | R | 1 |
|  |  | *msrA*, *msrD* | R | 1 |
|  |  | ***acrB* R717Q** | R | 1 (12) |
| Penicillins | Ampicillin (AMP) | ***bla*_TEM-1_** | R | 1 (31) |
|  |  | *bla*_OXA-1_, *bla*_OXA-7_ | R | 1 |
|  |  | Determinants of CEP and MEM | R | 2,3 |
| Cephalosporins | Extended-spectrum cephalosporins (CEP) | *bla*_CMY-2_ | R | 1 |
|  |  | *ampC* | R | 1 |
|  |  | ***bla*_CTX-M_** | R | 2 (37) |
|  |  | ***bla*_SHV_** | R | 2 (38) |
|  |  | *bla*_OXA-11_, *bla*_OXA-15_ | R | 2 |
|  |  | *bla*_TEM-10_, *bla*_TEM-124_ | R | 2 |
|  |  | Determinants of MEM | R | 3 |
| Carbapenems | Meropenem (MEM) | *bla*_AIM_ | R | 3 |
|  |  | *bla*_BIC_ | R | 3 |
|  |  | *bla*_DIM_ | R | 3 |
|  |  | *bla*_GES_ | R | 3 |
|  |  | *bla*_GIM_ | R | 3 |
|  |  | *bla*_IMI_ | R | 3 |
|  |  | *bla*_IMP_ | R | 3 |
|  |  | *bla*_KPC_ | R | 3 |
|  |  | *bla*_LMB_ | R | 3 |
|  |  | *bla*_NDM_ | R | 3 |
|  |  | *bla*_NMCA_ | R | 3 |
|  |  | *bla*_OXA-48_ like | R | 3 |
|  |  | *bla*_SIM_ | R | 3 |
|  |  | *bla*_SPM_ | R | 3 |
|  |  | *bla*_VIM_ | R | 3 |
| Polymixins | Colistin (CST) | *mcr-12345678* | R | 4 |

**Supplementary Table S8.** Distribution of ST assignments for 4389 public genomes as inferred by Pathogenwatch. The undetermined STs were explained by the absence or misassembly of one of the MLST genes (indicated with a question mark), suggesting that they are due to either sequencing or assembly errors. Novel STs are reported with a unique hash on the web application.

| **MLST ST** | **MLST Allelic Profile** | **No. of genomes (%)** | **Year range** |
| --- | --- | --- | --- |
| 1 | 1_1_1_1_1_1_5 | 2992 (68.17) | 1905-2019 |
| 2 | 1_1_2_1_1_1_5 | 1275 (29.05) | 1905-2017 |
| 2209 | 1_1_2_1_484_1_5 | 29 (0.66) | 1999-2015 |
| 2233 | 1_478_2_1_1_1_5 | 26 (0.59) | 2003-2016 |
| 2254 | 1_1_340_1_1_486_5 | 12 (0.27) | 2005-2012 |
| 2173 | 1_1_2_1_1_476_5 | 10 (0.23) | 2009-2017 |
| 8 | 1_1_2_3_1_1_5 | 5 (0.11) | 1936-2012 |
| 2244 | 1_1_2_616_1_1_5 | 4 (0.09) | 2006-2010 |
| 3803 | 1_1_2_1_1_1_733 | 4 (0.09) | 2009-2010 |
| 3677 | 1_1_2_1_1_665_5 | 3 (0.07) | 1996-2015 |
| 2230 | 1_1_1_1_1_1_526 | 3 (0.07) | 1986 |
| 2218 | 511_1_2_1_1_1_5 | 2 (0.05) | 2008-2010 |
| 2138 | 1_465_2_1_1_1_5 | 1 (0.02) | 2002 |
| 2160 | 1_1_385_3_1_1_5 | 1 (0.02) | 2011 |
| 2350 | 531_1_1_1_1_1_5 | 1 (0.02) | 2006 |
| 2352 | 532_1_2_3_1_1_5 | 1 (0.02) | 2005 |
| 2353 | 1_1_2_1_1_1_544 | 1 (0.02) | 2002 |
| 2355 | 1_1_2_1_1_1_500 | 1 (0.02) | 1998 |
| 2356 | 1_1_2_635_1_1_5 | 1 (0.02) | 1999 |
| 2360 | 1_1_1_1_1_498_5 | 1 (0.02) | 2010 |
| 3467 | 1_1_2_1_1_1_686 | 1 (0.02) | 2012 |
| 3476 | 1_1_2_1_1_1_687 | 1 (0.02) | 2013 |
| 3477 | 1_1_2_1_1_636_5 | 1 (0.02) | 2012 |
| 3738 | 1_1_2_880_1_1_5 | 1 (0.02) | 2016 |
| 3739 | 698_1_2_1_1_1_5 | 1 (0.02) | 1991 |
| 3891 | 1_1_1_910_1_1_5 | 1 (0.02) | 2015 |
| 2231 | 512_1_2_1_1_1_5 | 1 (0.02) | 1984 |
| 2331 | 1_1_1_633_1_1_5 | 1 (0.02) | 1973 |
| 2359 | 1_1_1_1_1_1_545 | 1 (0.02) | 1905 |
| Novel1 | 1_1_1_1_novel_1_5 | 1 (0.02) | 2016 |
| Novel2 | 1_1_2_novel_1_1_5 | 1 (0.02) | 2016 |
| Undetermined | 1_1_?_1_1_1_5 | 2 (0.05) | 2013 |
| Undetermined | 1_1_1_?_1_1_5 | 1 (0.02) | 2003 |
| Undetermined | 1_1_2_?_1_1_5 | 1 (0.02) | 2003 |
| Undetermined | ?_1_1_1_1_1_5 | 1 (0.02) | 1996 |

**Supplementary Table S9.** Distribution of known fluoroquinolone (Flq) resistance determinants (genes and mutations) in the 4389 public genomes in Pathogenwatch, and the linked genomic prediction of ciprofloxacin resistance provided by in the Antibiotics table.

| No. of Genomes | No. of Flq resistance genes | No. of Flq resistance mutations | Flq resistance determinants  identified in the genomes | Genomic prediction of ciprofloxacin resistance |
| --- | --- | --- | --- | --- |
| 1939 | 0 | 0 | None | Susceptible |
| 1219 | 0 | 1 | *gyrA*_S83F | Intermediate |
| 421 | 0 | 1 | *gyrA*_S83Y | Intermediate |
| 80 | 0 | 1 | *gyrB*_S464F | Intermediate |
| 68 | 0 | 1 | *gyrA*_D87N | Intermediate |
| 26 | 0 | 1 | *gyrA*_D87G | Intermediate |
| 16 | 0 | 1 | *gyrB*_S464Y | Intermediate |
| 14 | 0 | 1 | *gyrA*_D87Y | Intermediate |
| 1 | 0 | 1 | *gyrA*_D87V | Intermediate |
| 1 | 0 | 1 | *gyrA*_D87A | Intermediate |
| 1 | 0 | 1 | *gyrB*_Q465L | Intermediate |
| 158 | 0 | 2 | *gyrA*_S83F,*parE*_D420N | Intermediate |
| 27 | 0 | 2 | *gyrA*_S83F,*parC*_E84G | Intermediate |
| 23 | 0 | 2 | *gyrA*_S83F,*parE*_L416F | Intermediate |
| 7 | 0 | 2 | *gyrA*_S83F,*parC*_E84K | Intermediate |
| 4 | 0 | 2 | *gyrA*_S83F,*parC*_S80I | Intermediate |
| 3 | 0 | 2 | *gyrA*_S83Y,*parE*_D420N | Intermediate |
| 1 | 0 | 2 | *gyrB*_S464F,*gyrB*_Q465L | Intermediate |
| 1 | 0 | 2 | *gyrA*_D87N,*gyrB*_Q465L | Intermediate |
| 192 | 0 | 3 | *gyrA*_S83F,*gyrA*_D87N,*parC*_S80I | Resistant |
| 9 | 0 | 3 | *gyrA*_S83F,*gyrA*_D87G,*parC*_E84K | Intermediate |
| 6 | 0 | 3 | *gyrA*_S83F,*gyrB*_Q465R,*parE*_D420N | Intermediate |
| 5 | 0 | 3 | *gyrA*_S83F,*gyrA*_D87V,*parC*_S80I | Intermediate |
| 2 | 0 | 3 | *gyrA*_S83F,*gyrA*_D87G,*parC*_S80I | Intermediate |
| 1 | 0 | 3 | *gyrA*_S83F,*gyrA*_D87N,*parC*_E84K | Intermediate |
| 1 | 0 | 3 | *gyrA*_S83Y,*gyrA*_D87G,*parC*_S80I | Intermediate |
| 1 | 0 | 3 | *gyrA*_S83F,*gyrA*_D87G,*parE*_D420N | Intermediate |
| 1 | 0 | 3 | *gyrA*_S83F,*gyrA*_D87G,*parC*_E84G | Intermediate |
| 3 | 1 | 0 | *qnrS* | Intermediate |
| 87 | 1 | 1 | *qnrS*,*gyrA*_S83F | Resistant |
| 64 | 1 | 1 | *qnrS*,*gyrA*_S83Y | Resistant |
| 2 | 1 | 1 | *qnrB*,*gyrA*_S83F | Resistant |
| 2 | 1 | 2 | *qnrB*,*gyrA*_S83F,*parC*_E84K | Resistant |
| 3 | 1 | 3 | *qnrB*,*gyrA*_S83F,*gyrA*_D87N,*parC*_S80I | Resistant |
