## Additional File 4 Supplementary Figures for "A global resource for genomic predictions of antimicrobial resistance and surveillance of *Salmonella* Typhi at Pathogenwatch"

**Additional File 3. Supplementary Figures.**

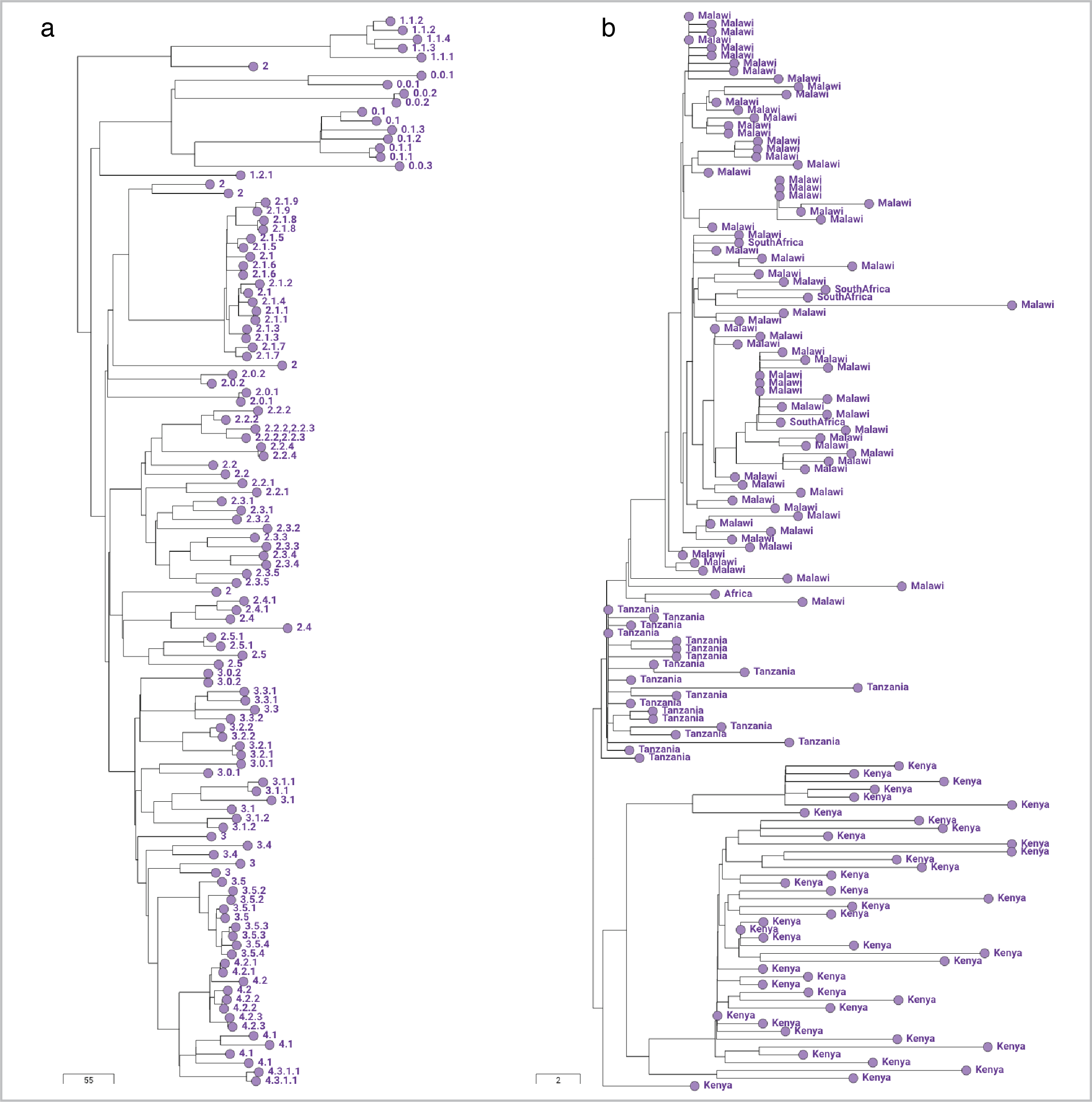

**Supplementary Figure S1. a** Pathogenwatch collection tree of 118 diverse *S.* Typhi genomes (dataset I) representing 62 genotypes (tree labels). **b** Pathogenwatch collection tree of 138 closely-related genomes from a clonal expansion within East Africa (dataset II). The tree nodes are labelled by country of isolation. The scale bars represent the number of SNP differences.

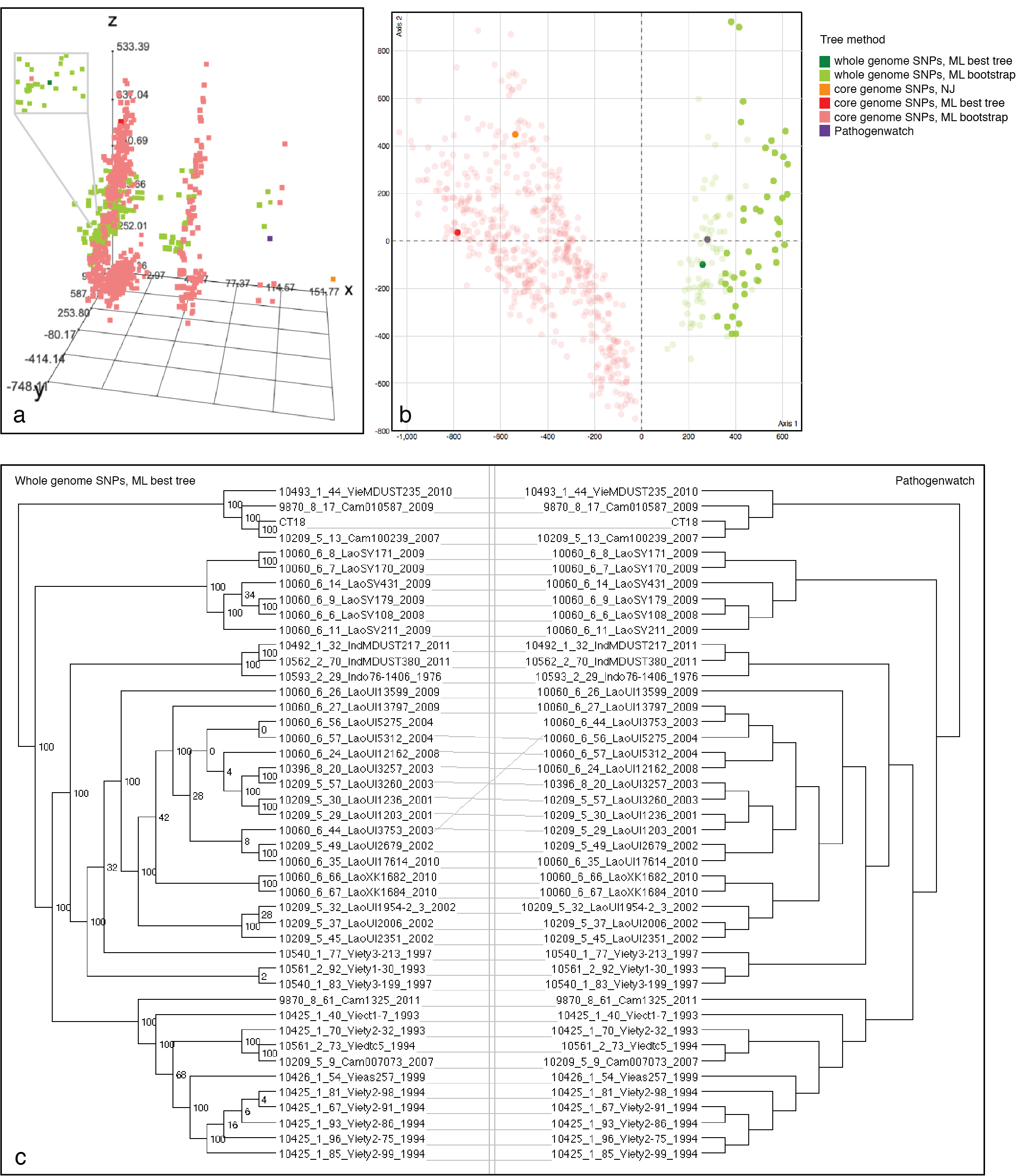

**Supplementary Figure S2.** Treespace of trees of **a** dataset I (118 genomes) and **b** dataset II (138 genomes) generated with four methods: 1) Pathogenwatch (purple); 2) Maximum likelihood on 9112 SNPs identified on an alignment of 3996 concatenated core genes estimated with roary (red); 3) Neighbour joining on the same alignment in 2) (orange); and 4) Maximum likelihood on 8598 SNPs called on a CT18-guided alignment (green). Five hundred bootstrap replicates were computed for each of the methods 2 (pink) and 4 (lime green). The inset unveils the position of the tree from method 4 (green). **c** Tanglegram comparison of the Pathogenwatch tree of 43 genomes from subclade 3.2.1 (right) with the CT18-guided alignment tree from method 4 (left).

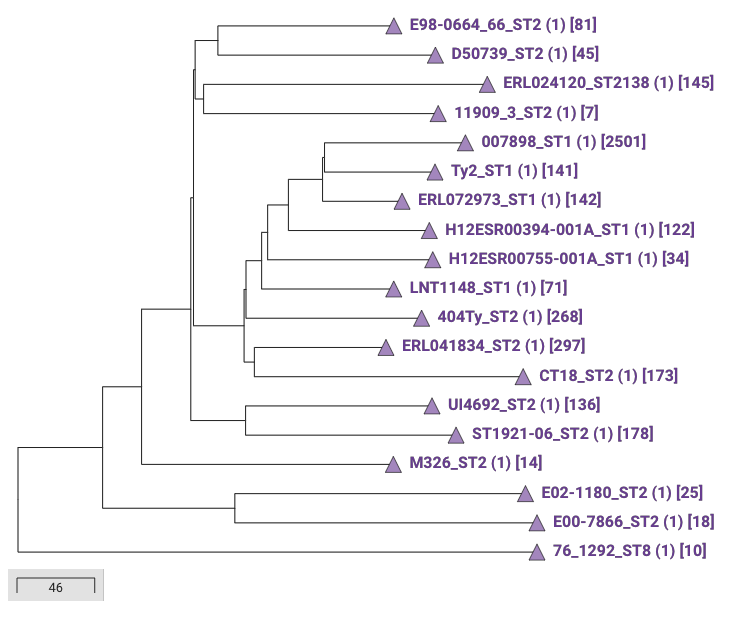

**Supplementary Figure S3.** The Typhi Pathogenwatch population Tree of 19 reference genomes, inferred from 2409 polymorphic sites found in 1639 of the 3916 core gene families. The numbers in parenthesis indicate the number of user genomes that have been sub-clustered with each reference in the tree. The numbers in brackets indicate the number of public genomes available on each subtree, which are also clustered with the user genomes.

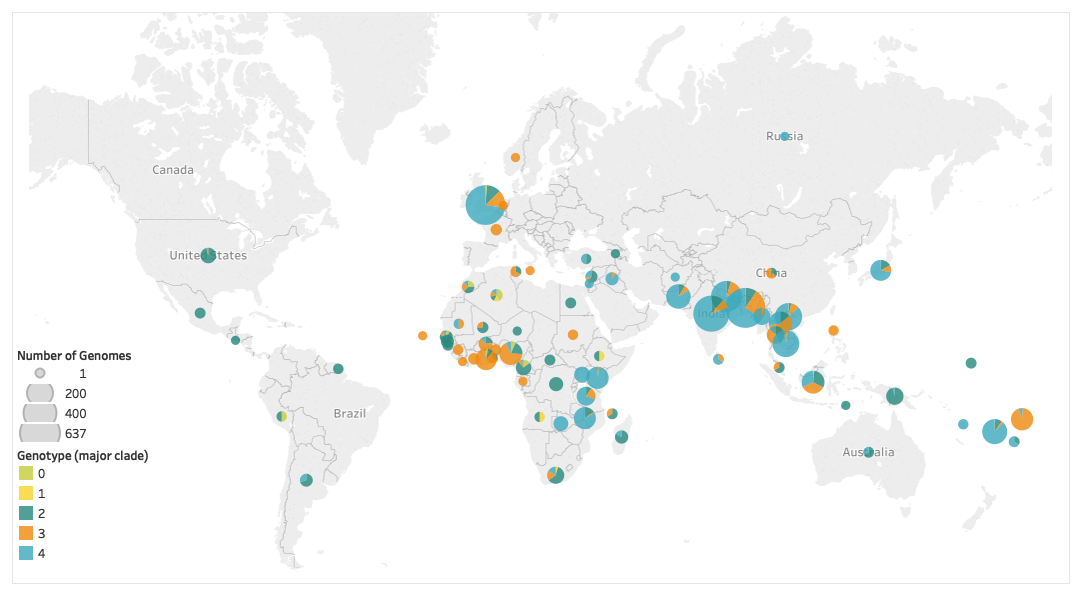
**Supplementary Figure S4.** Geographic distribution of public *S.* Typhi genomes available in Pathogenwatch. The size of the markers is proportional to the number of genomes in each country. The pie charts show the relative abundance of the four major clades per country. The visualization was created with Tableau Desktop v2018.3.

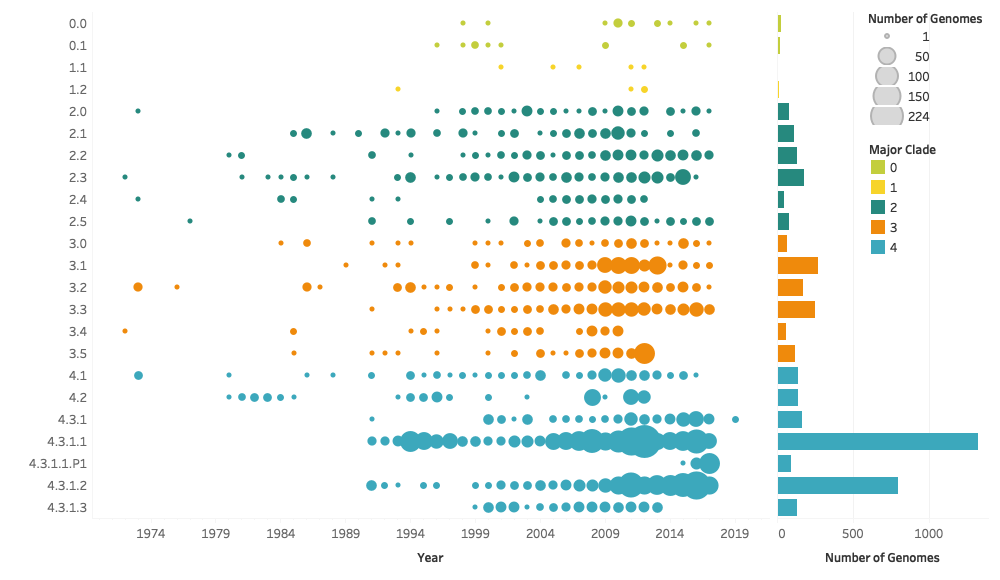
**Supplementary Figure S5.** Temporal distribution (left) and frequency (right) of the genotypes in the Typhi Pathogenwatch public collection of genomes. Genotypes were grouped by clade (e.g. 4.2) with the exception of 4.3, which was further discriminated into subclades. The markers on the timeline were coloured by the major clade they belong to, and their size is proportional to the number of genomes. The bar chart show the frequency of the genotypes, coloured by the major clade they belong to. Isolates collected before 1970 were excluded from the graphs. The visualization was created with Tableau Desktop v2018.3

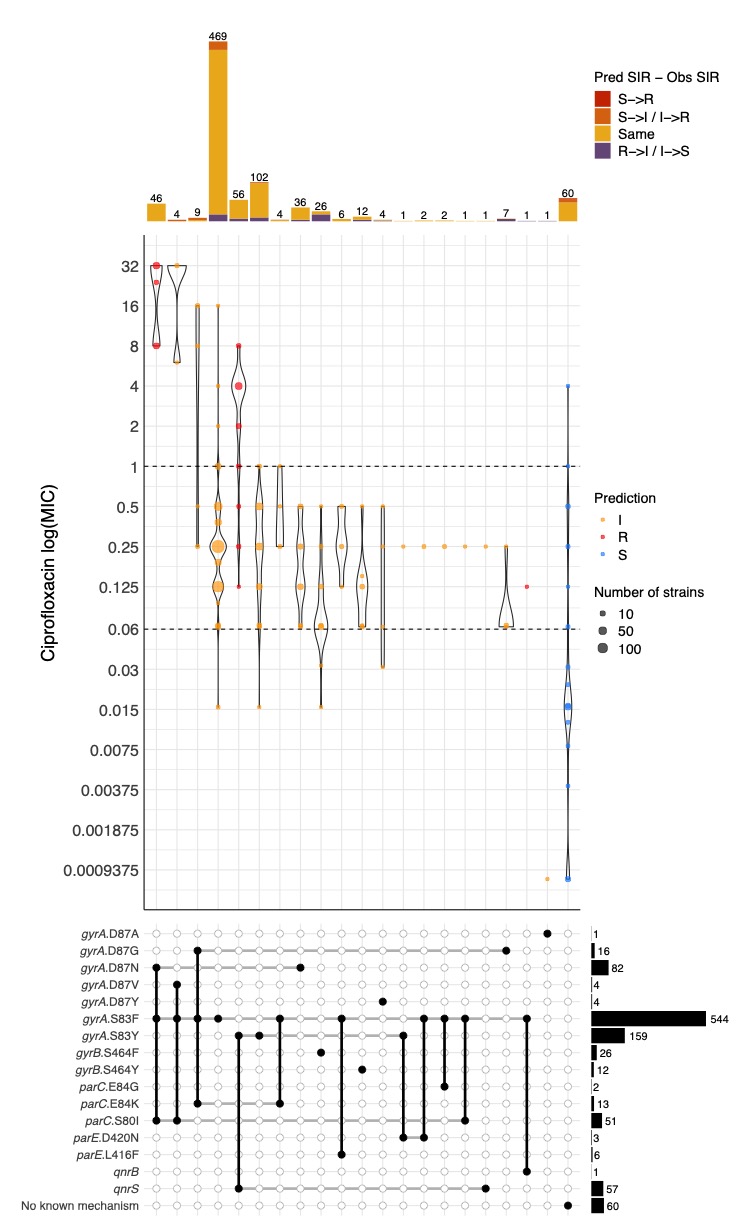

**Supplementary Figure S6.** Distribution of minimum inhibitory concentration (MIC) values (mg/L) for ciprofloxacin in a collection of 889 *S.* Typhi isolates with different combinations of genetic mechanisms that are known to confer resistance to this antibiotic. Dashed horizontal lines on the violin plots mark the CLSI clinical breakpoint for ciprofloxacin. Point colours inside violins represent the genotypic AMR prediction by Pathogenwatch on each combination of mechanisms. Bar plots on the top show the abundance of genomes with each combination of mechanisms. Bar colours represent the differences between the predicted and the observed SIR (i.e. red for a predicted susceptible mechanism when the observed phenotype is resistant).

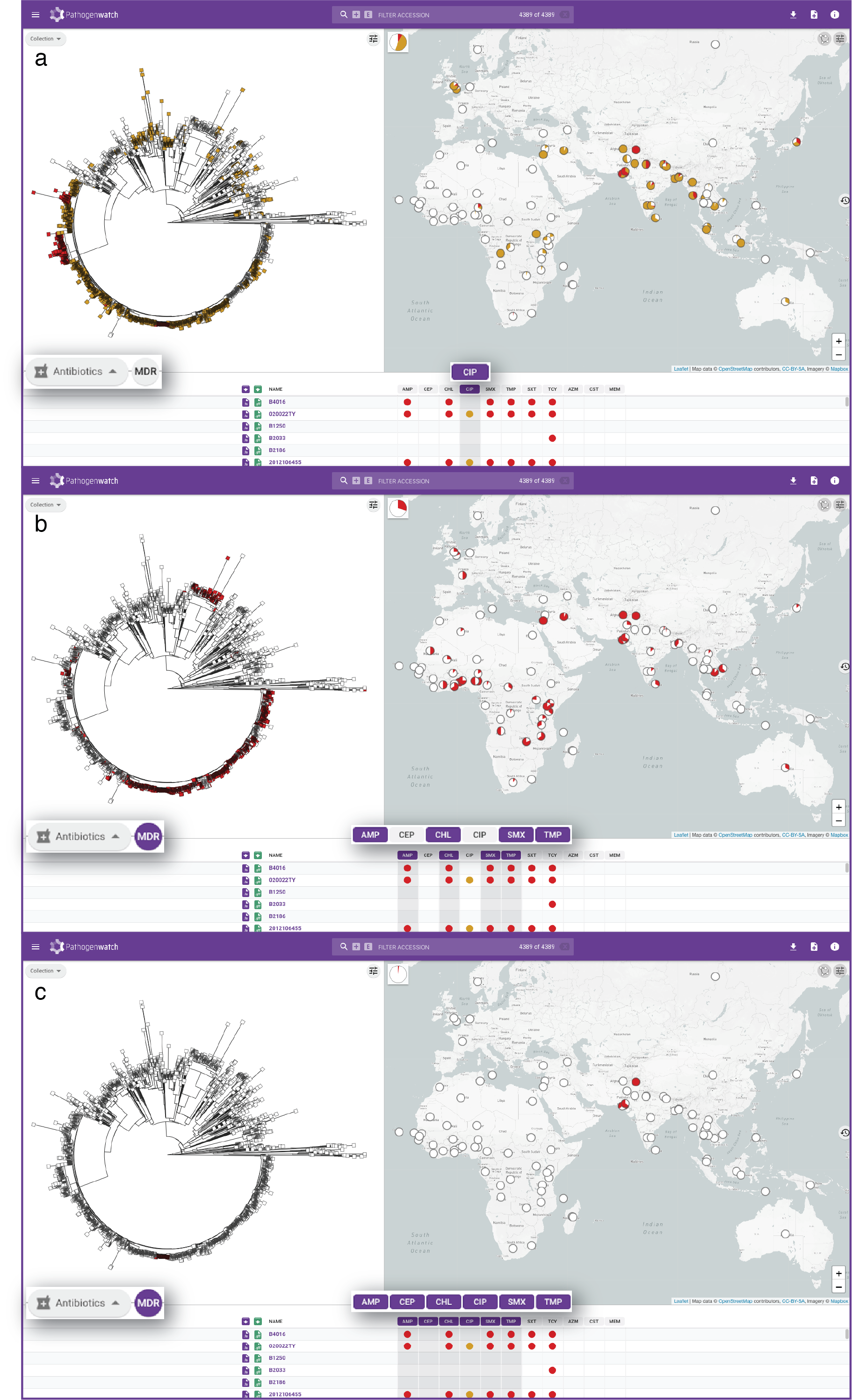

**Supplementary Figure S7.** Distribution of predicted ciprofloxacin (CIP)-resistant (**a**), multi-drug resistant (**b**), and extremely-drug resistant (**c**) *S.* Typhi on the map and tree of 4389 genomes. Tree nodes are coloured red (resistant), yellow (intermediate), or white (no resistance predicted). The map pie charts display the proportion of resistant/intermediate genomes at each location. The expandable pie chart at the top-left corner of the map view indicates the proportion of resistant/intermediate genomes for the whole set of 4389 genomes.

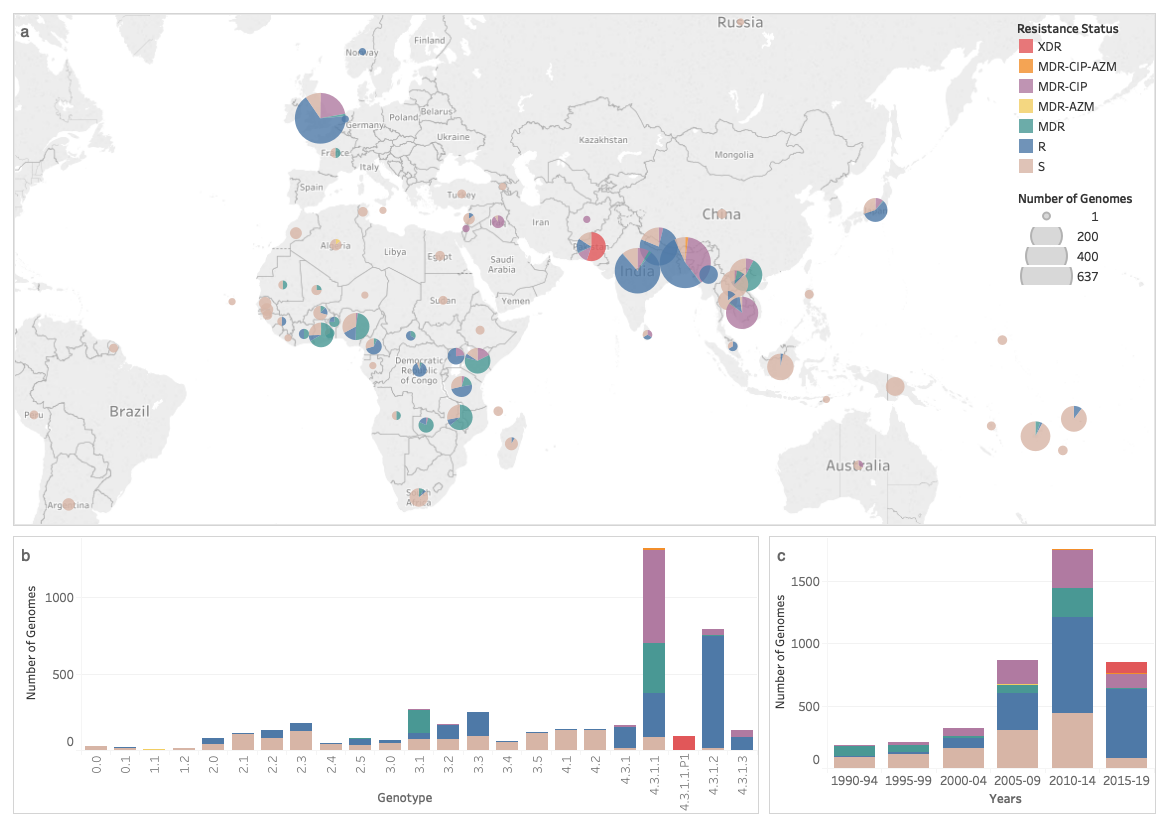

**Supplementary Figure S8.** Distribution of the Pathogenwatch-predicted resistance profiles for the public genomes by country (a), genotype (b) and year (c). Isolates collected before 1970 were excluded from (c). MDR: resistant to ampicillin, chloramphenicol, sulfamethoxazole, and trimethoprim. XDR: resistant to ampicillin, chloramphenicol, sulfamethoxazole, trimethoprim, ciprofloxacin, and extended-spectrum cephalosporins. MDR-CIP|AZM: MDR with additional resistance to ciprofloxacin and or azithromycin. R: resistant to at least one antibiotic but not classified in the previous categories. S: no resistance found. The visualization was created with Tableau Desktop v2018.3.

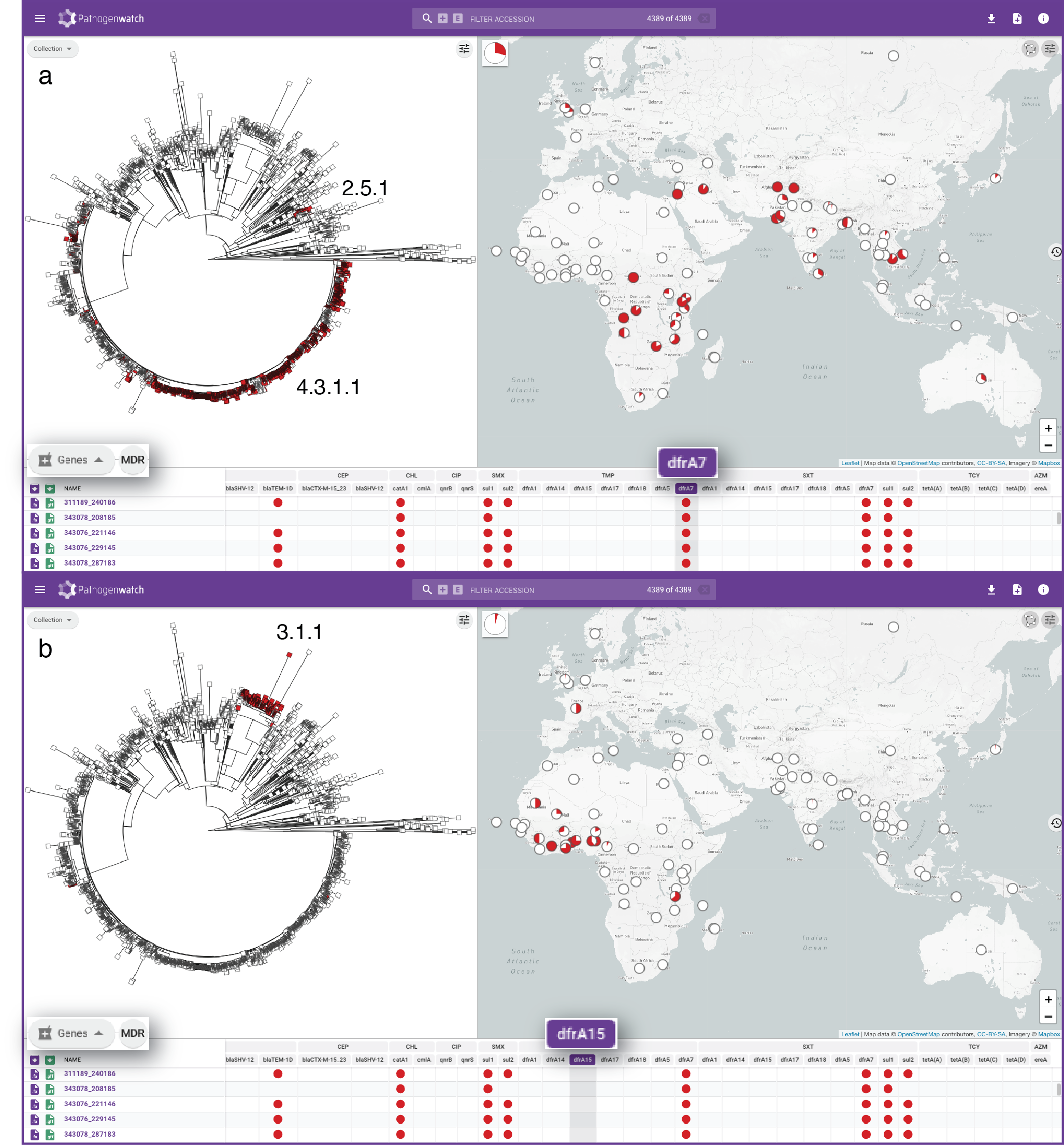

**Supplementary Figure S9.** Distribution of trimethoprim resistance genes *dfrA7* (**a**) and *dfrA15* (**b**). Genes are grouped by antibiotic on the genes table. Top panel: widespread distribution of *dfrA7* in Asia and East Africa, largely linked to subclades 4.3.1.1 and 2.5.1. Bottom panel: localised distribution of *dfrA15* in West Africa associated with subclade 3.3.1.

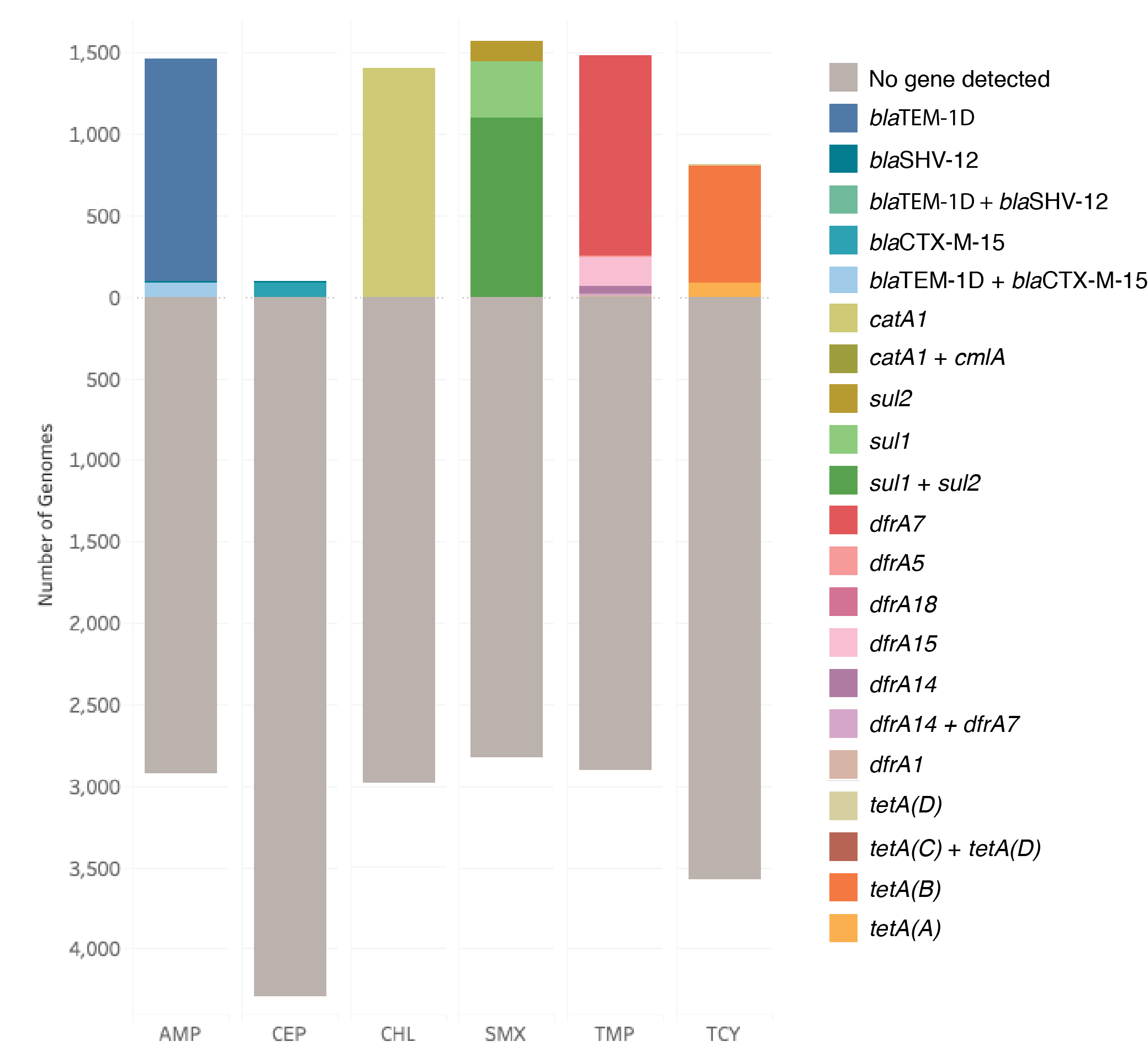
**Supplementary Figure S10.** Distribution of genes in the Typhi Pathogenwatch AMR library linked to resistance to ampicillin (AMP), extended-spectrum cephalosporins (CEP), chloramphenicol (CHL), sulfamethoxazole (SXT), trimethoprim (TMP) and tetracycline (TCY) across the public genomes. Genomes predicted to be resistant to each antibiotic are shown above the 0 on the y-axis, and those without any known gene found below the 0. The visualization was created with Tableau Desktop v2018.3.

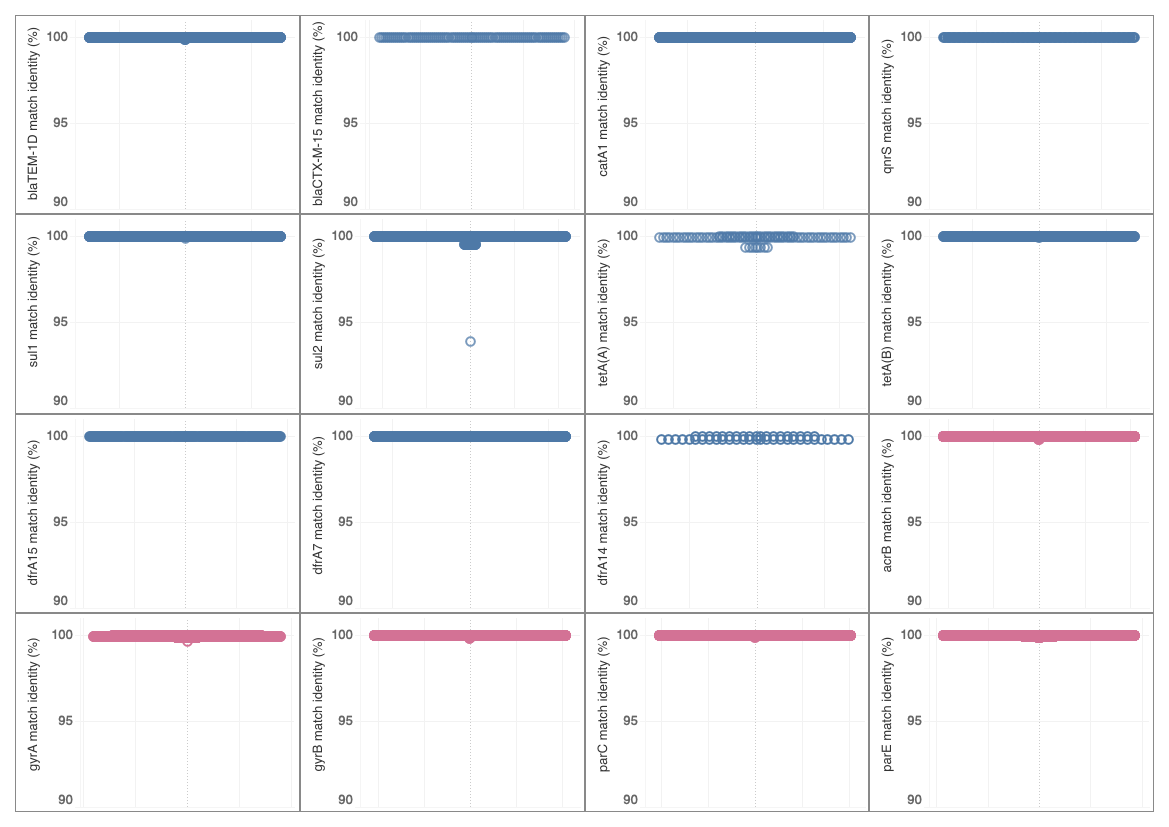

**Supplementary Figure S11.** Distribution of identity values of the blast matches identified in the 4389 public genomes to the genes in the AMR library. Only genes with at least 10 matches in the public genomes are shown. Blue circles: acquired resistance genes. Pink circles: housekeeping genes target of resistance mutations. The visualization was created with Tableau Desktop v2018.3.

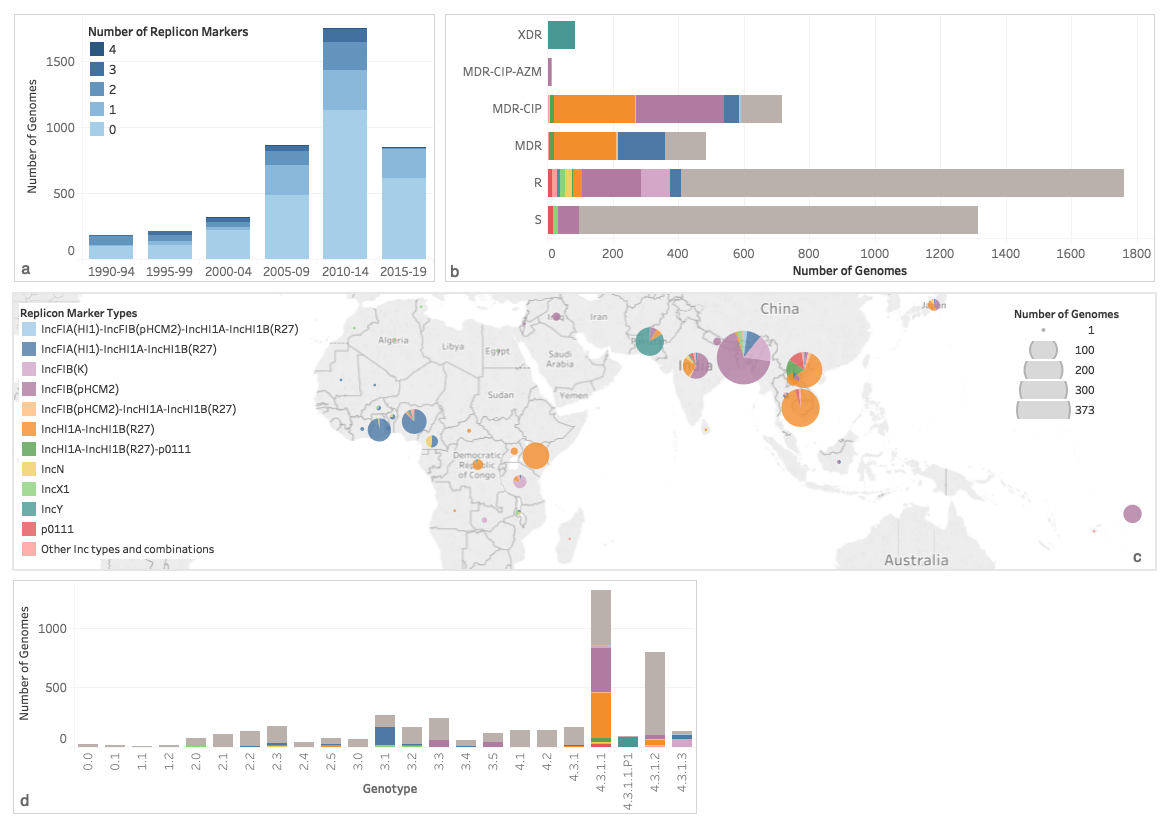

**Supplementary Figure S12.** Distribution of plasmid replicon marker types across the 4389 public genomes in Typhi Pathogenwatch. **a** Number of Replicon markers per year interval. The genomes from isolates collected before 1990 were excluded. **b** Distribution of replicon marker types across the different predicted resistance profiles grouped as in Supplementary Figure 4. **c** Geographical distribution. **d** Distribution of replicon types across the different predicted genotypes. The visualization was created with Tableau Desktop v2018.3.

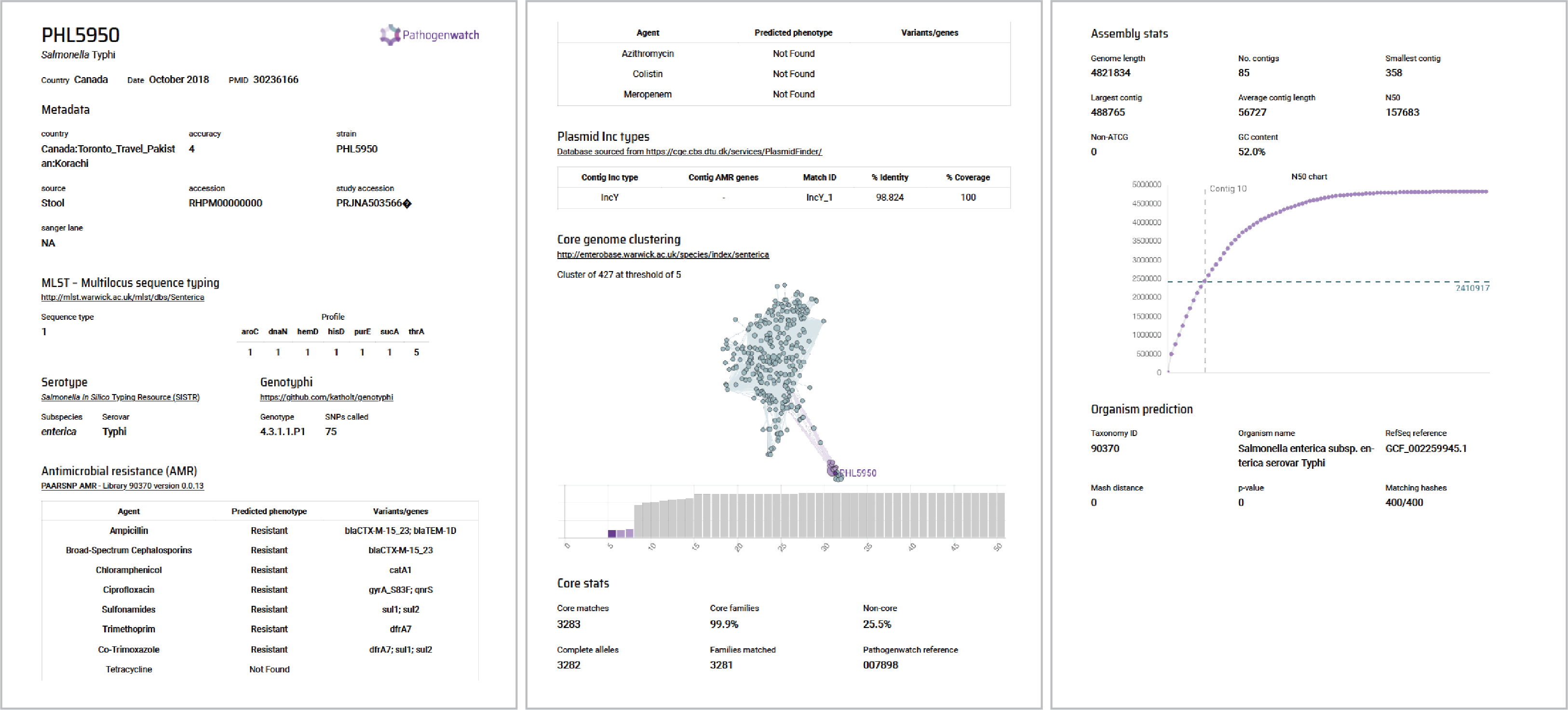

**Supplementary Figure S13.** Pathogenwatch genome report for assembly PHL595.
